# Pulmonary mRNA-LNP Vaccines for Rapid and Durable Protection Against Bacterial Infection

**DOI:** 10.64898/2025.12.19.695407

**Authors:** Xiaoli Wei, Anqi Wei, Yu Miao, Zhou Yuan, Guanghui Li, Dexuan Lei, Yinyu Ma, Zhiwei Guo, Yuansong Sun, Tianhao Ding, Kaisong Tian, Qin He, Zui Zhang, Lianfeng Fan, Changyou Zhan

## Abstract

Pulmonary bacterial infections remain a major clinical challenge. Although vaccination reduces infection rates and mortality, the vulnerable post-vaccination immunity gap can still result in infection and vaccine failure. In addition, effective vaccines are unavailable for many clinically important bacterial pathogens. Here, we report a pulmonary mRNA-lipid nanoparticle (mRNA-LNP) vaccine incorporating a novel ionizable lipid engineered for localized high-level expression, which elicits both rapid and durable protections against bacterial lung infections, effectively bridging this critical window of vulnerability. Intratracheal delivery of mRNA-LNP rapidly primes lung neutrophils and macrophages into a transcriptionally pre-activated state, enhancing their phagocytic activity and enabling rapid, antigen-independent bacterial clearance during the early post-vaccination period (approximately 1-7 days). Subsequently, vaccination induces potent antigen-specific adaptive responses, conferring sustained protection against both laboratory and clinical drug-resistant *Pseudomonas aeruginosa* strains. Single-cell transcriptomics and immune profiling reveal coordinated activation of innate and adaptive immune programs. This dual-phase immune response exemplifies a paradigm-shifting vaccine design that integrates innate and adaptive immunity to confer both immediate and long-term protection. Our findings establish a mechanistic basis for rapid antibacterial defense and highlight pulmonary mRNA-LNP vaccination as a promising strategy for combating respiratory infections.

---

Bacterial infections continue to pose a significant global health burden, particularly in the context of increasing antibiotic resistance. Pulmonary infections such as pneumonia, caused by pathogens including *Streptococcus pneumoniae*, *Pseudomonas aeruginosa*, and *Acinetobacter baumannii*, account for millions of deaths annually and remain among the leading causes of mortality in vulnerable populations, including children, the elderly, and immunocompromised individuals^1,2^. Hospital-acquired pneumonia (HAP), which develops 48 hours or more after hospital admission, is one of the most common healthcare-associated infections and is associated with high morbidity and mortality^3–5^. The growing prevalence of multidrug-resistant (MDR) strains has further limited therapeutic options and underscores the urgent need for effective prophylactic strategies^6,7^.

Vaccines represent a powerful tool in preventing bacterial infections and reducing antimicrobial use. Although current vaccines such as pneumococcal conjugate vaccines have achieved notable clinical success, effective vaccines against many clinical important respiratory pathogens are still lacking^8,9^. While the development of bacterial vaccines is hindered by the intrinsic complexity of antigen selection and pathogen variability, the immunogenicity and protective efficacy of such vaccines are also heavily dependent on the design and performance of adjuvants, underscoring the need for innovative delivery systems and immune-potentiating strategies. Furthermore, following immunization, the generation of pathogen-specific adaptive immunity typically requires several days to weeks, creating a post-vaccination immunity gap during which the host remains susceptible to infection ^10–12^. This is particularly problematic in hospital environments, where patients-and even accompanying individuals-can be exposed to pathogenic bacteria within hours to days after admission^13^. As a result, pre-hospitalization vaccination may not provide timely protection in this scenario. Therefore, there is a critical need to develop next-generation vaccines capable of inducing rapid immune protection to bridge this vulnerability window. Such vaccines would be particularly valuable in acute exposure settings or among high-risk populations, where immediate defense against pathogens is essential for effective infection control^14^.

Recent advances in nanotechnology-enabled mRNA delivery systems, particularly lipid nanoparticles (LNPs), have revolutionized vaccine development by enabling efficient cytosolic delivery and robust antigen expression^15,16^. Beyond serving as delivery vehicles, mRNA-LNPs also possess inherent adjuvant properties^17^, as both the mRNA, which can activate pattern recognition receptors (PRRs)^18,19^, and the lipid components, which promote robust T follicular helper (Tfh) cell responses^20^, contribute to innate immune activation and enhanced antigen presentation^21^, thereby amplifying adaptive immunity. While most of these advancements have focused on systemic administration, especially intramuscular injection, the potential of LNPs for mucosal immunization via the pulmonary route has attracted growing interest due to the lungs’ rich immune network and large surface area^22,23^. Delivering mRNA-LNP vaccines directly to the lungs may offer unique advantages in combating respiratory infections, including localized immune activation, rapid innate response engagement, and mucosal protection^24–26^. While current studies primarily focus on aerosol stability and delivery efficiency^27,28^, the immunological mechanisms and local immune modulation in the lung following mRNA-LNP vaccine administration are less well understood.

In this study, we developed an mRNA-LNP vaccine incorporating a novel ionizable lipid specifically engineered for localized high-level expression and evaluated its capacity to confer both immediate and long-term protection against bacterial pulmonary infections following intrapulmonary immunization. Through comprehensive *in vivo* evaluation in murine models, we demonstrate that pulmonary mRNA-LNP vaccination leads to rapid immune activation, reduction in early bacterial burden, and robust adaptive immune responses. Mechanistically, we performed single-cell RNA sequencing and immune cell profiling to dissect the early innate responses and downstream adaptive pathways triggered by localized antigen expression. Our findings suggest a promising vaccination strategy to close the immunity gap, offering both rapid protection against acute bacterial challenges and durable immunity for long-term defense.

## Results

### Optimization of LNPs for localized high expression

The ionizable lipid plays a central role in mRNA encapsulation and delivery, critically influencing both the *in vivo* protein expression efficiency and adjuvant properties of the vaccine^20,29,30^. First, we systematically designed and synthesized a series of ionizable lipids by incorporating malonate (MA1 - 13) and cyclohexane derivatives (CY1 - 13) as structure skeletons, chosen for their synthetic versatility and tunable modification properties (Supplementary Fig. S1a)^31,32^. These cationic lipids were then formulated into LNPs encapsulating luciferase (Luc) mRNA and evaluated for their size (Supplementary Fig. S1b), encapsulation efficiency (Supplementary Fig. S1c) and *in vivo* transfection ability after intramuscular injection (i.m.) and intratracheal nebulization (i.t.) (Supplementary Fig. S1d-g). We found that lipids bearing a four-branched tail achieved superior mRNA expression, while reducing the branch length markedly diminished transfection efficiency, highlighting the importance of both lipid branching and sufficient hydrophobicity. Notably, cyclohexane-core lipids (CY series) promoted preferential expression at the injection site, with significantly reduced hepatic expression compared to MA lipids and the FDA-approved SM102. Among the candidates, CY7-incorporating a 4-dimethylaminopiperidine head group (Fig.1a) emerged as the lead compound, exhibiting the strongest *in vivo* expression and a favorable biodistribution profile, with consistently higher expression at the injection site relative to the liver (Fig.1b-e). To further characterize tissue-selective expression, Cre mRNA-loaded LNPs were fabricated (Supplementary Fig. S2) and administered intratracheally to Lox-3xSTOP-Lox-tdTomato (Ai9) reporter mice^33^. Three days post-delivery, lung and liver tissues were harvested for fluorescence imaging. Histological analysis confirmed that LNP_CY7_ induced robust tdTomato expression in the lungs, whereas no detectable fluorescence was observed in the liver, in stark contrast to LNP_SM102_ (Fig. 1f and Supplementary Fig. S3), highlighting CY7’s selective expression at the site of administration.

**Fig 1.**
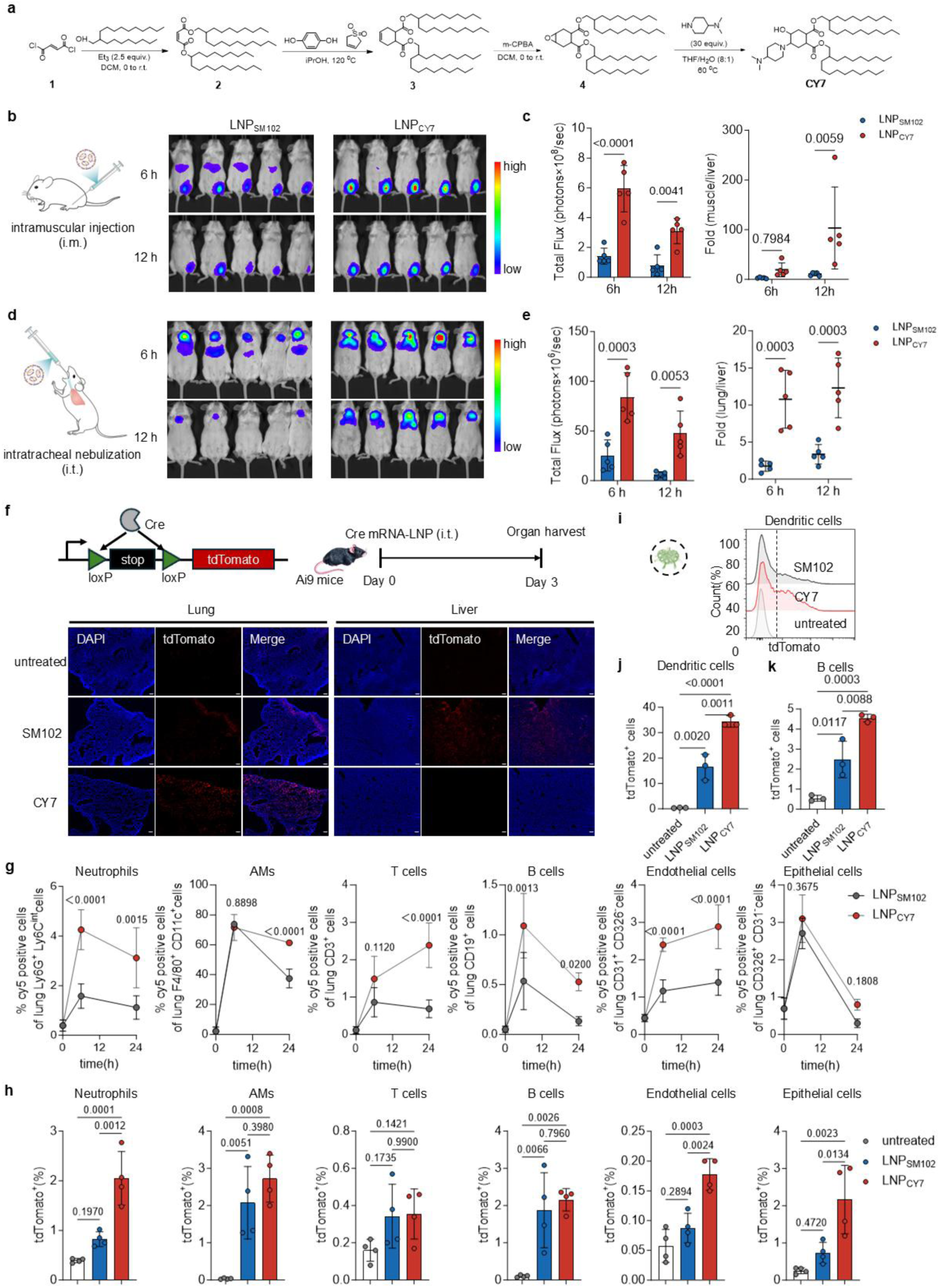
LNP_CY7_ demonstrated high localized mRNA expression. **a,** Synthetic route of the ionizable lipid CY7. **b,** Whole-body bioluminescence imaging of mice treated with Luc mRNA-LNPs following intramuscular (i.m.) injection. **c,** Quantification of total bioluminescence flux at the injection site after i.m. administration, and fold change in bioluminescence at the site of administration (muscle) relative to the liver (n=5). **d,** Whole-body bioluminescence imaging of mice treated with Luc mRNA-LNPs following intratracheal (i.t.) nebulization. **e,** Quantification of total bioluminescence flux in the lung after i.t. nebulization, and fold change in bioluminescence at the site of administration (lung) relative to the liver (n=5). **f,** Fluorescence imaging of lung and liver tissue sections from Ai9 reporter mice 3 days after intratracheal administration of Cre mRNA-LNPs formulated with CY7 or SM102. Scale bar, 100 µm. **g,** Cellular uptake of LNPs by distinct pulmonary cell populations, assessed at 6 and 24 hours after intratracheal administration of Cy5-labeled mRNA-LNPs (n=4). **h,** Expression of tdTomato in distinct pulmonary cell populations of Ai9 mice at 72 hours post intratracheal delivery of Cre mRNA-LNPs (n=4). **i,** Representative flow plots of tdTomato^+^ dendritic cells in the mediastinal lymph nodes. **j and k,** Quantification of tdTomato^+^ dendritic cells (j) and B cells (k) in the mediastinal lymph nodes (n=3). Data represent mean ± s.d. Statistical significance was determined using one-way or two-way ANOVA with Tukey’s multiple comparisons test in GraphPad Prism 10.

To assess cell-type specific uptake and expression within the lung, flow cytometry was performed following intratracheal nebulization. Alveolar macrophages represented the primary population taking up fluorescent dye-labeled mRNA-loaded LNPs, and LNP_CY7_ demonstrated consistently higher uptake across all major pulmonary cell types compared with LNP_SM102_ (Fig. 1g and Supplementary Fig. S4). Correspondingly, LNP_CY7_ elicited elevated Cre mRNA expression in neutrophils, endothelial cells, and epithelial cells relative to LNP_SM102_ (Fig. 1h). Beyond the lung, enhanced mRNA expression was also detected in the draining lymph node. LNP_CY7_ induced markedly stronger expression in antigen presenting cells, including dendritic cells and B cells, compared with LNP_SM102_ (Fig. 1i–k). Collectively, these results demonstrate that LNP_CY7_ enables more efficient mRNA delivery and expression in key immune cell types within the respiratory tract and draining lymph nodes, underscoring its potential as a promising platform for pulmonary mRNA vaccination.

### Pulmonary delivery of mRNA-LNPs offers rapid protection against bacterial lung infection

To investigate the immune responses following pulmonary vaccination, we selected PcrV-a key component of the type Ⅲ secretion system of *Pseudomonas aeruginosa* as a model antigen and prepared mRNA-LNP formulations using either CY7 or SM102 as the ionizable lipid via a microfluidic mixing system ^34^ (Supplementary Fig. S5a and b). Dynamic light scattering analysis showed that the resulting LNPs had an Z-average diameter of approximately 75 nm (Supplementary Fig. S5c), with both formulations exhibiting encapsulation efficiencies exceeding 95% (Supplementary Fig. S5d). Cryo-electron microscopy further revealed that the LNPs exhibited uniform and well-defined spherical morphology, with no observable aggregation and clearly resolved nanostructures (Fig. 2a). We then performed a dose-escalation study to determine the optimal pulmonary dose (Supplementary Fig. S6). Administration of 5 μg of mRNA in the mRNA-LNP formulation led to significant body weight loss and even mortality, along with pronounced acute lung inflammation and focal accumulation of inflammatory cells 24 hours post-administration. In contrast, mice that received 1 μg or 2 μg of mRNA, like the control group, showed no obvious inflammation, tissue damage, or changes in body weight. Importantly, the 2 μg dose elicited higher PcrV-specific antibody titers than the 1 μg dose (Supplementary Fig. S7) and was therefore chosen as the optimal dose for subsequent vaccination experiments.

**Fig 2.**
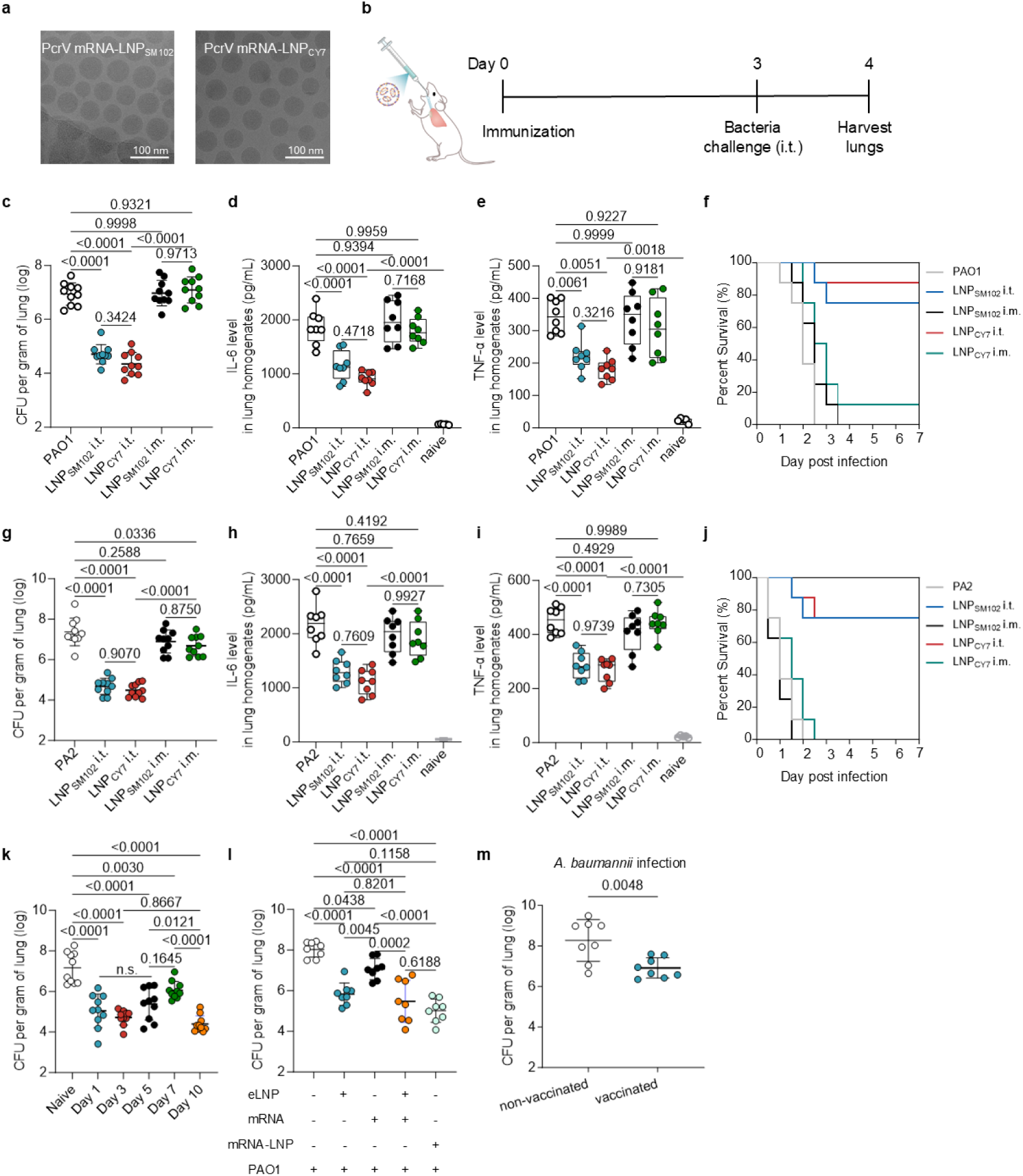
Pulmonary delivery of mRNA-LNPs offers rapid protection against bacterial lung infection. **a,** Representative cryo-EM images of PcrV mRNA-LNPs formulated with SM102 or CY7 ionizable lipids. Scale bar = 100 nm. **b,** Schematic representation of LNP administration and subsequent bacterial challenge. **c,** Bacterial burden in the lungs of *P. aeruginosa* (PAO1)-infected mice. Mice were immunized via intratracheal nebulization or intramuscular injection of PcrV mRNA-LNPs, followed by intrapulmonary bacterial challenge three days later. At 24 hours post-infection, lungs were collected, homogenated and plated for Colony-Forming Unit (CFU) enumeration (n=10). **d, e,** IL-6 (d) and TNF-α (e) concentrations in lung homogenates collected from PAO1 infected mice across different treatment groups (n=8). **f,** Survival curve of mice after pulmonary challenge with PAO1 (n=8). **g,** Bacterial burden in the lungs of mice infected with the clinical *P*. aeruginosa strain PA2 (n=10). **h, i,** IL-6 (h) and TNF-α (i) concentrations in lung homogenates collected from PA2-infected mice across different treatment groups (n=8). **j,** Survival curve of mice after pulmonary challenge with PA2 (n=8). **k,** Bacterial enumeration in PAO1-infected lungs measured 24 hours after pulmonary infection on days 1, 3, 5, 7, and 10 post-vaccination (with a booster administered on day 7) (n=10). **l,** Mice received a single administration with PcrV mRNA-LNP or eLNP or mRNA or eLNP mixed with mRNA and challenged with PAO1 three days later. Lungs were collected at 24 h post-infection and plated for CFU enumeration (n=8). **m,** Bacterial enumeration in the lungs of mRNA-LNP_CY7_ immunized mice infected with *Acinetobacter baumannii* (n=8). Data represent mean ± s.d. Statistical significance was determined using one-way ANOVA with Tukey’s multiple comparisons test in GraphPad Prism 10.

Surprisingly, when mice were immunized via intratracheal administration and challenged with the *Pseudomonas aeruginosa* PAO1 strain three days later, we unexpectedly observed that both mRNA-LNP_CY7_ and mRNA-LNP_SM102_ conferred substantial protection. Despite the short interval between vaccination and infection, the pulmonary bacterial burden was markedly reduced at 24 hours post-challenge (Fig. 2b and c). Notably, this effect was not observed following intramuscular immunization. This unexpected early protection was further supported by significantly decreased levels of the pro-inflammatory cytokines IL-6 and TNF-α (Fig. 2d and e). Under high-dose bacterial challenge, pulmonary vaccination substantially improved survival outcomes (Fig. 2f), demonstrating robust short-term protection. In contrast, intramuscular administration failed to elicit early protection. Similar protective trends were observed against a clinical *P. aeruginosa* isolate designated PA2, a carbapenem-resistant strain isolated from a sputum sample (Fig. 2g-j). These findings highlight the unique potential of pulmonary mRNA-LNP vaccination to rapidly control bacterial lung infections, bridging the early immunity gap prior to the establishment of adaptive responses.

Next, we investigated the duration of this protective effect (Fig. 2k). We found that the short-term protection persisted on day 1, 3, and 5 post-vaccinations. Although this protection partially declined by day 7, administration of a booster dose on day 7 restored protection. We then explored the components responsible for the observed protective immune effect (Fig. 2l). Administration of empty LNP_CY7_ (eLNPs) at an equivalent lipid dose to the mRNA-LNP group significantly reduced the bacterial burden in the lung. Although this reduction was slightly less pronounced than that observed in the mRNA-LNP group, the difference was not statistically significant. Co-administration of free PcrV mRNA with empty LNPs significantly reduced lung bacterial burden to levels comparable to those achieved with the fully formulated mRNA-LNP vaccine. These results indicate that the LNP carrier is not only essential for efficient mRNA delivery but also plays a critical role in mediating short-term protective immune responses.

To determine whether the observed protection was antigen-specific, we first measured antibody responses three days post-vaccination. As expected within this early window, no detectable levels of antigen-specific IgA, IgM, or IgG were found (data not shown), suggesting that adaptive humoral immunity was not yet established. To further probe the nature of the protection, mice vaccinated with PcrV mRNA-LNPs were intratracheally infected with *A. baumannii*, a pathogen that does not express PcrV. Interestingly, pulmonary bacterial burden was significantly reduced (Fig. 2m), indicating the presence of antigen-independent innate immune protection.

### LNP-induced priming of neutrophils and macrophages drives early antibacterial defense

Given the central role of neutrophils and macrophages in innate immunity and bacterial clearance, their phagocytic capacity was first examined. Following pulmonary administration of the mRNA-LNP vaccine, both neutrophils and macrophages displayed significantly enhanced bacterial uptake, indicating that these cells were primed for heightened antimicrobial activity (Fig. 3a-d). Notably, although neutrophil recruitment to the lung increased transiently shortly after vaccination, it returned to baseline within 24 hours, indicating that the early protective effect was not simply due to an elevated number of neutrophils and macrophages at the time of bacterial challenge (Fig. 3e and f). To further explore additional mechanisms contributing to rapid protection, we assessed neutrophil extracellular trap (NET) formation in the lung after bacterial infection using immunofluorescence staining for myeloperoxidase (MPO) and citrullinated histone H3 (CitH3). Vaccination alone did not induce detectable NETs (Supplementary Fig. S8). In contrast, following bacterial infection, vaccinated mice exhibited markedly increased NET formation compared with controls, indicating that pulmonary mRNA-LNP immunization primes neutrophils to also undergo enhanced NETosis upon pathogen encounter, thereby contributing to early antibacterial protection (Fig. 3g and Supplementary Fig. S9). Next, neutrophils and macrophages were selectively depleted using anti-Ly6G antibody and clodronate liposomes, respectively, to directly assess their contribution^35–38^. Depletion of either cell type markedly impaired the antibacterial efficacy of the mRNA-LNP vaccine, confirming their essential role in early protection (Fig. 3h-i and Supplementary Fig. S10). Collectively, these findings demonstrate that pulmonary administration of mRNA-LNPs primes neutrophils and macrophages into a pre-activated state, thereby enhancing their phagocytic and bactericidal functions.

**Fig 3.**
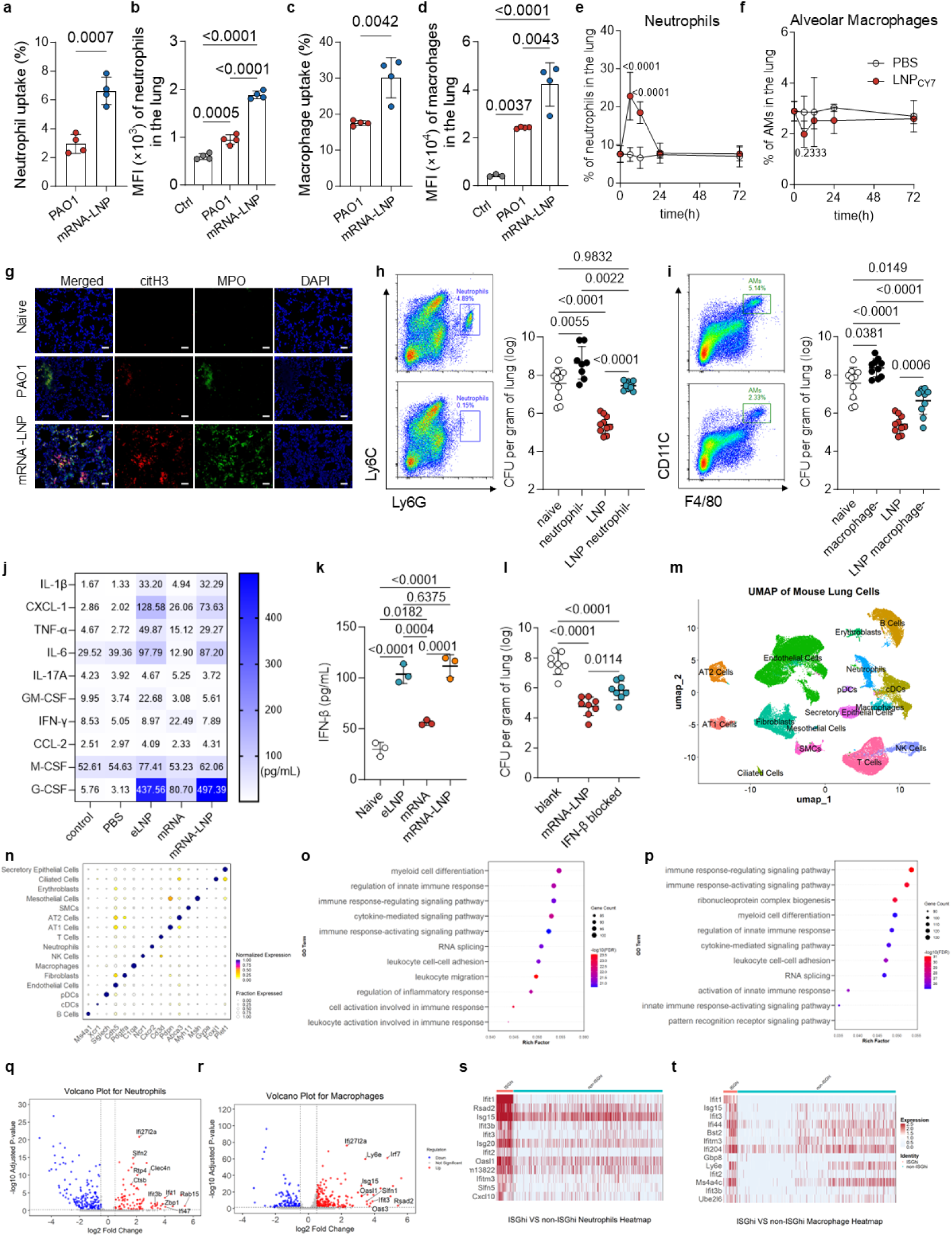
LNP-induced priming of neutrophils and macrophages drives early antibacterial defense. **a, b,** Cellular uptake percentage (a) and median fluorescent intensity (MFI) (b) of DiI-labeled PAO1 bacteria by lung neutrophils 3 days post-immunization, quantified using flow cytometry (n=4). **c, d,** Cellular uptake percentage (c) and MFI (d) of DiI-labeled PAO1 bacteria by lung macrophages 3 days post-immunization, quantified using flow cytometry (n=4). **e, f,** Lung neutrophil (e) and alveolar macrophages (f) proportion at indicated post-vaccination time points (n=4). **g,** Representative immunofluorescence images of NET formation in the lungs of PAO1-infected mice and PAO1-infected vaccinated mice, 24 hours after intratracheal bacterial challenge. Staining shows citrullinated histone H3 (Cit-H3, red), myeloperoxidase (MPO, green), and nuclei (DAPI, blue). Scale bar, 50 µm. **h,** Flow cytometry showing depletion of lung neutrophils by anti-Ly6G antibody, and bacterial colony counts in the lungs with or without neutrophil depletion (n=8-10). **i,** Flow cytometry showing depletion of lung alveolar macrophages by clodronate liposome, and bacterial colony counts in the lungs with or without alveolar macrophage depletion (n=10). **j,** Cytokine and chemokine levels in BLAF quantified 6 hours post-vaccination using a flow cytometry-based multiplex panel (n=3). **k,** IFN-β levels in BALF of vaccinated mice 6 hours post-vaccination, measured by ELISA (n=3). **l,** Bacterial enumeration in the lungs of infected mice with or without IFN-β blockade. Mice were intratracheally administered PcrV mRNA-LNP with or without IFN-β antibody blockade, followed by intrapulmonary bacterial challenge three days later. Lung homogenates were collected 24 hours post-infection and plated for CFU enumeration (n =8). **m,** Uniform manifold approximation and projection representation of scRNA-seq data annotated with cell types. **n,** Dot plot showing the identified cell types and their representative marker genes based on scRNA-seq analysis. Dot size indicates the proportion of cells within each cluster expressing the given marker gene, while dot color reflects the min-max normalized average expression level of each gene in the corresponding cell type. **o, p,** GO analysis of neutrophils (o) and macrophages (p) 72 hours post pulmonary mRNA-LNP vaccination. **q, r,** Volcano plot showing differentially expressed genes (DEGs) in neutrophils (q) and macrophages (r) from control and mRNA-LNP-treated lungs. The x-axis represents the log2 fold change of gene expression, and the y-axis represents the -log10 of the adjusted P-values. **s, t,** Heatmap of the top differentially expressed genes (DEGs) between ISG^hi^ and non-ISG^hi^ neutrophils (s) and macrophages (t) identified from single-cell RNA sequencing data. Data represent mean ± s.d. Statistical significance was determined using one-way or two-way ANOVA with Tukey’s multiple comparisons test in GraphPad Prism 10.

To better characterize the local immune milieu associated with neutrophil and macrophage pre-activation, we analyzed a panel of cytokines and chemokines in bronchoalveolar lavage fluid (BALF) (Fig. 3j-k). Levels of Interleukin-6 (IL-6), C-X-C motif chemokine ligand 1 (CXCL-1), Granulocyte Colony-Stimulating Factor (G-CSF) and Interferon beta (IFN-β) were markedly elevated after mRNA-LNP administration. IL-6 is a well-characterized inflammatory mediator known to promote dendritic cell activation, thereby facilitating the development of subsequent adaptive immune responses^20,39^. Temporal profiling revealed that IL-6 induction was transient, returning to baseline within 24 hours (Supplementary Fig. S11a). Both CXCL-1 and G-CSF contribute to neutrophil recruitment and functional priming, with G-CSF further enhancing phagocytic and bactericidal activity^40,41^. Furthermore, pulmonary delivery of mRNA-LNPs resulted in a transient elevation of IFN-β in bronchoalveolar lavage fluid, suggesting engagement of the type I interferon response (Supplementary Fig. S11b). Co-administration of a neutralizing anti-IFN-β antibody alongside PcrV mRNA-LNP partially diminished the protective effect, as evidenced by a significant increase in lung bacterial burden relative to the mRNA-LNP-only group (Fig. 3l). Upon bacterial infection, IFN-I signaling is known to support bacterial clearance not only by enhancing phagocytic and bactericidal activity, but also by promoting neutrophil extracellular trap formation^42–44^. Importantly, the transient elevation of these factors appeared to be driven primarily by the LNP components themselves, highlighting the intrinsic adjuvant properties of the lipid nanoparticles. Together, these findings not only reinforce the adjuvant effect of eLNP in potentiating antigen-specific immunity but also provide a mechanistic explanation for the rapid, short-term protective immunity conferred by pulmonary mRNA-LNP vaccination.

Although levels of inflammatory cytokines returned to baseline within a short period, neutrophils and macrophages still exhibited enhanced phagocytic activity. To explore the underlying mechanisms, we performed single-cell RNA sequencing (scRNA-seq) on mouse lung tissues 72 hours post-immunization. Uniform Manifold Approximation and Projection (UMAP) analysis identified 15 transcriptionally distinct clusters, encompassing structural parenchymal cells as well as diverse immune populations (Fig. 3m and n). Gene Ontology (GO) and Reactome pathway analysis of neutrophils and macrophages revealed enrichment of multiple innate immune signaling pathways, including immune response-activating signaling pathway and cytokine-mediated signaling pathway, indicating that mRNA-LNP exposure induces innate immune reprogramming (Fig. 3o-p and Supplementary Fig. S12). This reprogramming triggers sustained downstream transcriptional programs, allowing phagocytes to maintain enhanced functionality even after the transient inflammatory response had waned. Volcano plot analysis further demonstrated significant upregulation of canonical interferon-stimulated genes (ISGs) such as Ifi27l2a, Rtp4, Ifit1, and Irf7 in neutrophils and macrophages at 72 hours (Fig. 3q and r)^45,46^. Sub-clustering identified a transcriptionally distinct population enriched for ISG expression, which we designated ISG^hi^ (Fig. 3s and t). Cells within this cluster exhibited a robust type I IFN transcriptional signature, previously implicated in promoting antibacterial defense in specific contexts^47,48^.

Taken together, these findings suggest that pulmonary mRNA-LNP administration places neutrophils and macrophages in a primed functional state, accompanied by transient inflammatory signaling and sustained transcriptional changes. This immune configuration enables enhanced phagocytic activity and provides rapid, short-term antibacterial protection prior to the establishment of adaptive immunity.

### LNP_CY7_ enhances adjuvant activity for adaptive immune activation

Building on the confirmed activation of innate immunity and its resulting short-term protection, we next evaluated the adjuvant properties of LNP in orchestrating long-term adaptive immune responses. mRNA encoding the *P. aeruginosa* antigen PcrV was encapsulated into LNP_CY7_ or LNP_SM102_ and delivered via either intratracheal or intramuscular administration. Draining lymph nodes were harvested at defined time points post-vaccination to assess local IL-6 production, a cytokine known to facilitate T follicular helper (Tfh) cell differentiation and germinal center (GC) formation^49^ (Fig. 4a and b). Both delivery routes led to elevated IL-6 levels compared to naïve controls; LNP_CY7_ consistently induced significantly higher IL-6 production than SM102-formulated LNPs, highlighting its superior adjuvant potency.

**Fig 4.**
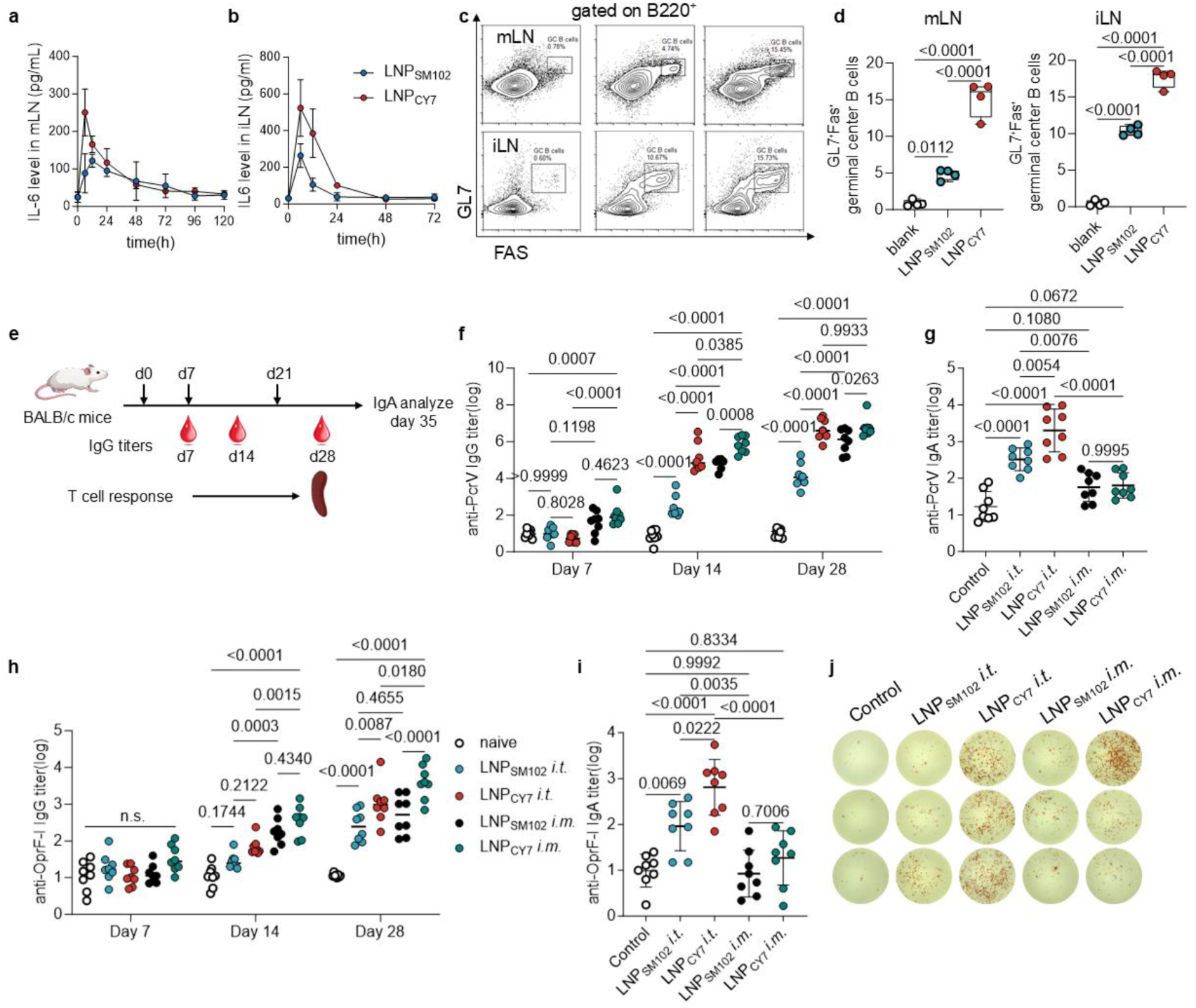
mRNA-LNP_CY7_ immunization elicited strong humoral and cellular immune responses. **a,b,** IL-6 levels in lymph nodes following immunization with PcrV loaded LNP_SM102_ or LNP_CY7_ via intramuscular or intratracheal administration. IL-6 levels were measured by ELISA in lysates of mediastinal lymph nodes (mLN) (a) or iliac lymph nodes(iLN) (b) at indicated time points post-immunization. **c, d,** Representative flow cytometry plots (c) and quantification (d) of GC B cells (B220^+^ FAS^+^GL7^+^) in draining lymph nodes on day 28 post-immunization via different administration routes. **e,** Schematic illustration of the prime-boost vaccination strategy and timeline for the analysis of immune responses. **f,** Serum IgG titers against PcrV in immunized mice at different time points post-immunization (n=8). **g,** Antigen-specific IgA titers against PcrV in bronchoalveolar lavage fluid (BALF) 35 days post-immunization (n=8). **h,** Serum IgG titers against OprF-I in immunized mice at different time points post-immunization (n=8). **i,** Antigen-specific IgA titers against OprF-I in BALF 35 days post-immunization (n=8). **j,** ELISpot plate images indicating IFN-γ-producing T cells in immunized mice. Data represent mean ± s.d. Statistical significance was determined using one-way or two-way ANOVA with Tukey’s multiple comparisons test in GraphPad Prism 10.

Next, germinal center (GC) responses in the draining lymph nodes were examined, as GCs serve as essential sites for B-cell proliferation, somatic hypermutation, and affinity maturation, thereby shaping the quality and durability of humoral immunity^50^. Flow cytometry revealed that LNP_CY7_ induced markedly greater GC B cell (FAS⁺GL7⁺) differentiation compared to SM102-based formulations following both administration routes (Fig. 4c and d), further supporting the enhanced immune adjuvant capacity of LNP_CY7_. To elucidate the mechanism underlying this enhanced immunogenicity, we compared the adjuvant properties of empty LNPs (eLNPs) co-administered with recombinant PcrV protein (rPcrV). Notably, eLNP_CY7_ and eLNP_SM102_ triggered comparable levels of inflammatory cytokines in the draining lymph nodes (Supplementary Fig. S13), suggesting that the enhanced adjuvant activity of mRNA-LNP_CY7_ for adaptive immunity is likely attributed to more efficient *in situ* antigen expression, rather than intrinsic differences in innate inflammatory signaling.

### mRNA-LNP_CY7_ immunization elicited robust humoral and cellular immune responses

Having established the superior adjuvant activity of LNP_CY7_ in promoting innate immune activation and germinal center formation, we next assessed whether these immunostimulatory properties translate into robust antigen-specific adaptive immune responses. To this end, we developed mRNA-LNP vaccines targeting *Pseudomonas aeruginosa*, incorporating two well-characterized protective antigens: the full-length PcrV protein and a fusion of outer membrane proteins OprF and OprI^51^. Corresponding mRNAs were synthesized and individually encapsulated into LNP using a microfluidic mixing platform. The two LNP formulations were then mixed and administered together at the same mRNA dose for immunization. The resulting mRNA-LNPs exhibited uniform particle sizes (∼75 nm) and high encapsulation efficiencies (>95%) (Supplementary Fig. S5 and S14).

To evaluate humoral immunity, BALB/c mice were immunized via either intratracheal or intramuscular routes, followed by booster doses (Fig. 4e). ELISA analysis showed that antibody responses remained low one week after the initial pulmonary administration but were robustly elevated following booster immunization against both PcrV and OprF-I. Notably, LNP_CY7_ outperformed SM102 across both routes of administration, inducing significantly higher levels of antigen-specific IgG (Fig. 4f and h). Isotype profiling further indicated a Th1/Th2-balanced or slightly Th2-biased response (Supplementary Fig. S15). To assess mucosal immunity, antigen-specific IgA levels in bronchoalveolar lavage fluid were quantified on day 35. Pulmonary administration of LNP elicited robust IgA responses in the lung, whereas intramuscular injection failed to induce detectable IgA, demonstrating the advantage of intrapulmonary vaccination in promoting localized mucosal responses. Moreover, CY7-based formulations generated higher IgA titers than SM102-based formulations, highlighting the superior immunogenicity of mRNA-LNP_CY7_ (Fig. 4g and i).

We further analyzed cellular immunity by isolating splenocytes one week after the final boost and stimulating them with PcrV and OprF-I antigens *ex vivo*. ELISpot assays showed significantly more IFN-γ producing cells in the LNP_CY7_ group than in the SM102 group (Fig. 4j and Supplementary Fig. S16). These results demonstrate that mRNA-LNP_CY7_ vaccination elicits strong and multifunctional adaptive immune responses, highlighting its potential as a next-generation platform for bacterial vaccines.

### mRNA-LNP_CY7_ elicits protective adaptive immune responses against pulmonary and systemic P. aeruginosa infections

To assess the protective efficacy of adaptive immune responses elicited by mRNA-LNP vaccination, mice were immunized with either a combination of PcrV mRNA-LNP_CY7_ and OprF-I mRNA-LNP_CY7_ or a combination of PcrV mRNA-LNP_SM102_ and OprF-I mRNA-LNP_SM102_ (Fig. 5a). Neither intrapulmonary nor intramuscular vaccination resulted in significant changes in body weight compared with control mice, indicating good tolerability (Fig. 5b). Vaccinated mice were subsequently challenged intratracheally with *Pseudomonas aeruginosa* strain PAO1 or a carbapenem-resistant clinical isolate (PA2). Both intratracheal nebulization and intramuscular immunization of mRNA-LNP_CY7_ significantly reduced lung bacterial burden compared with unvaccinated controls and the mRNA-LNP_SM102_ group, following challenge with either PAO1 or PA2 (Fig. 5c and g). Notably, although intramuscular vaccination also conferred measurable protection in the lung, which may rely on a small amount of systemically induced IgG reaching the lung mucosa either through FcRn-mediated transport or via passive transudation due to increased vascular permeability during infection^52,53^, intratracheal immunization consistently resulted in superior protective efficacy, likely owing to the induction of local mucosal immune responses.

**Fig 5.**
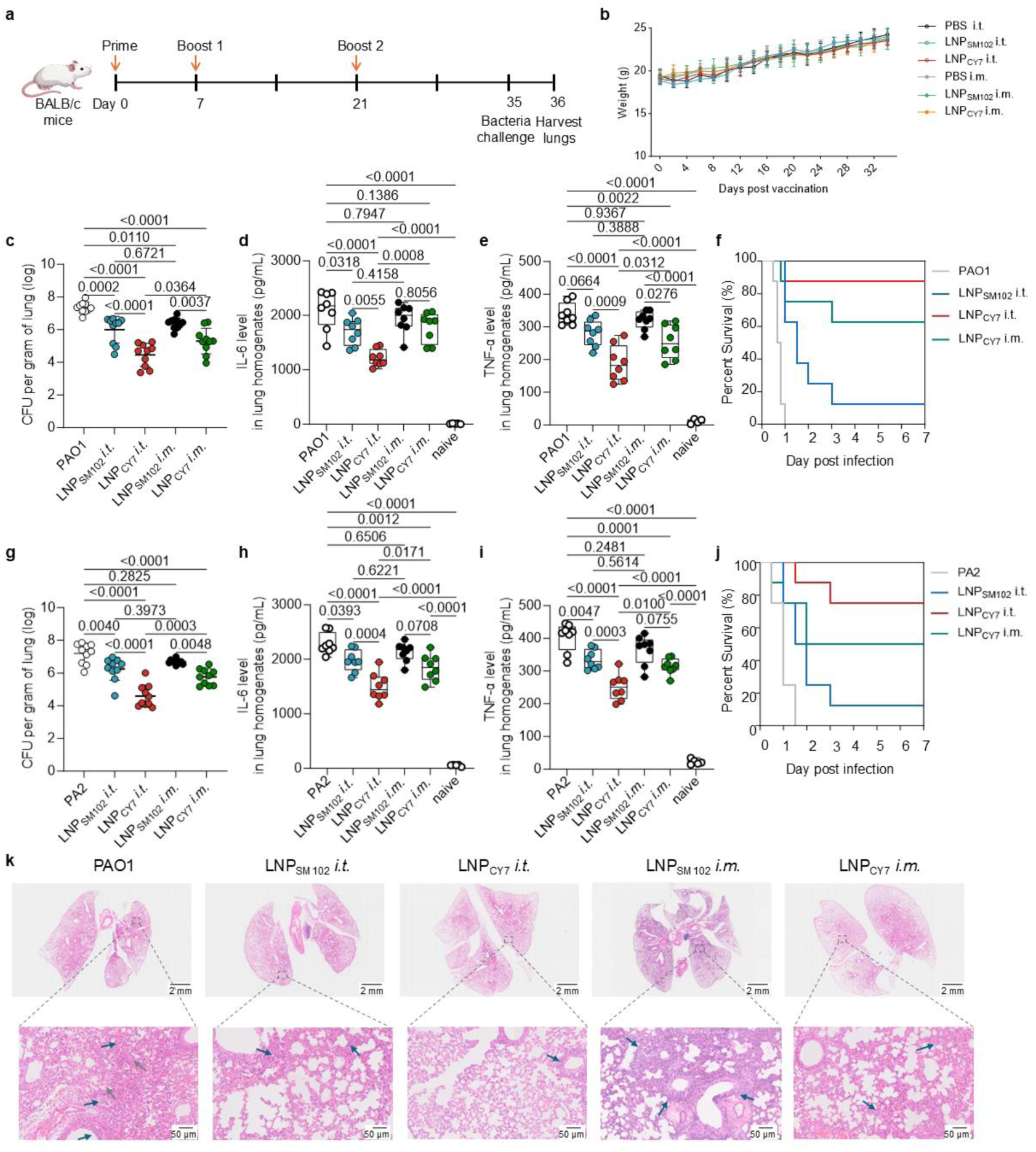
mRNA-LNP_CY7_ vaccination protects against *P. aeruginosa* infections in pulmonary infection model. **a,** Timeline of mouse immunization and subsequent *P. aeruginosa* challenge. **b,** Body weight monitoring following intratracheal and intramuscular administration of mRNA-LNPs. Control mice received PBS only. **c,** Lung bacterial burden in PAO1-infected mice. Lung homogenates were collected 24 hours post-infection and plated for bacterial enumeration (n=10). **d, e,** IL-6 (d) and TNF-α levels (e) in lung homogenates of PAO1-infected mice, quantified by ELISA (n=8). **f,** Survival curves of mice following pulmonary challenge with *P. aeruginosa* PAO1 strain (n=8). **g,** Lung bacterial burden in PA2-infected mice (n=10). **h, i,** IL-6 levels (h) and TNF-α levels (i) in the lung homogenates of PA2-infected mice, quantified by ELISA (n=8). **j,** Survival curves of mice following pulmonary challenge with PA2 (n=8). **k,** Representative hematoxylin and eosin (HE)-stained lung sections from PAO1-infected control and immunized mice. Blue arrows indicate diffuse neutrophilic infiltration; grey arrows indicate alveolar damage and hemorrhage. Scale bars: 2 mm (overview) and 50 µm (high magnification). Data represent mean ± s.d. Statistical significance was determined using one-way ANOVA with Tukey’s multiple comparisons test in GraphPad Prism 10.

Consistent with reduced bacterial burden, vaccination significantly alleviated lung inflammation, as evidenced by decreased levels of the proinflammatory cytokines IL-6 and TNF-α (Fig. 5d-e and h-i). Survival analysis demonstrated that mRNA-LNP_CY7_ vaccination significantly improved survival outcomes in mice with pulmonary *P. aeruginosa* infection (Fig. 5f and j). Notably, intratracheal immunization of mRNA-LNP_CY7_ conferred the highest survival benefit, outperforming both intratracheal LNP_SM102_ and intramuscular LNP_CY7_ immunization, highlighting the superior efficacy of mRNA-LNP_CY7_ delivered via the mucosal route. Further histopathological examination of lung tissues collected 24 hours post-infection revealed extensive pulmonary damage in unvaccinated control mice, including diffuse neutrophilic infiltration, alveolar collapse, and alveolar hemorrhage. In contrast, vaccinated groups exhibited markedly reduced inflammation and preservation of alveolar architecture, with the most pronounced protection observed in the mRNA-LNP_CY7_ intratracheal immunization group (Fig. 5k). Collectively, these results indicate that mRNA-LNP_CY7_ vaccination not only enhances bacterial clearance and improves survival but also effectively protects lung tissue from acute inflammatory injury, with intratracheal delivery providing superior mucosal immunity and overall protective efficacy.

Having demonstrated robust pulmonary protection, we next investigated whether mRNA-LNP_CY7_ could confer systemic immunity in a disseminated infection model. Mice immunized with mRNA-LNP vaccines were challenged intravenously with either PAO1 or PA2, and bacterial burdens in major organs-including the heart, lung, liver, spleen, and kidney-were quantified 24 hours post-infection. Compared to unvaccinated controls, both vaccine formulations significantly reduced bacterial loads in all tested organs, with mRNA-LNP_CY7_ conferring notably superior protection over mRNA-LNP_SM102_ against both laboratory and clinical strains, irrespective of whether vaccines were administered intramuscularly or via intratracheal nebulization (Fig. 6a-e, 6f-j). Survival curve analysis further demonstrated that vaccination with LNP_CY7_ significantly prolonged the survival of mice, regardless of the route of administration (Fig. 6k-l). In contrast, mice immunized with luciferase (Luc) mRNA-LNPs as a control showed no detectable protective effect (Supplementary Fig. S17). These findings indicate that the protection observed is antigen-specific and mRNA-LNP_CY7_-induced adaptive immunity provides robust systemic antibacterial defense.

**Fig 6.**
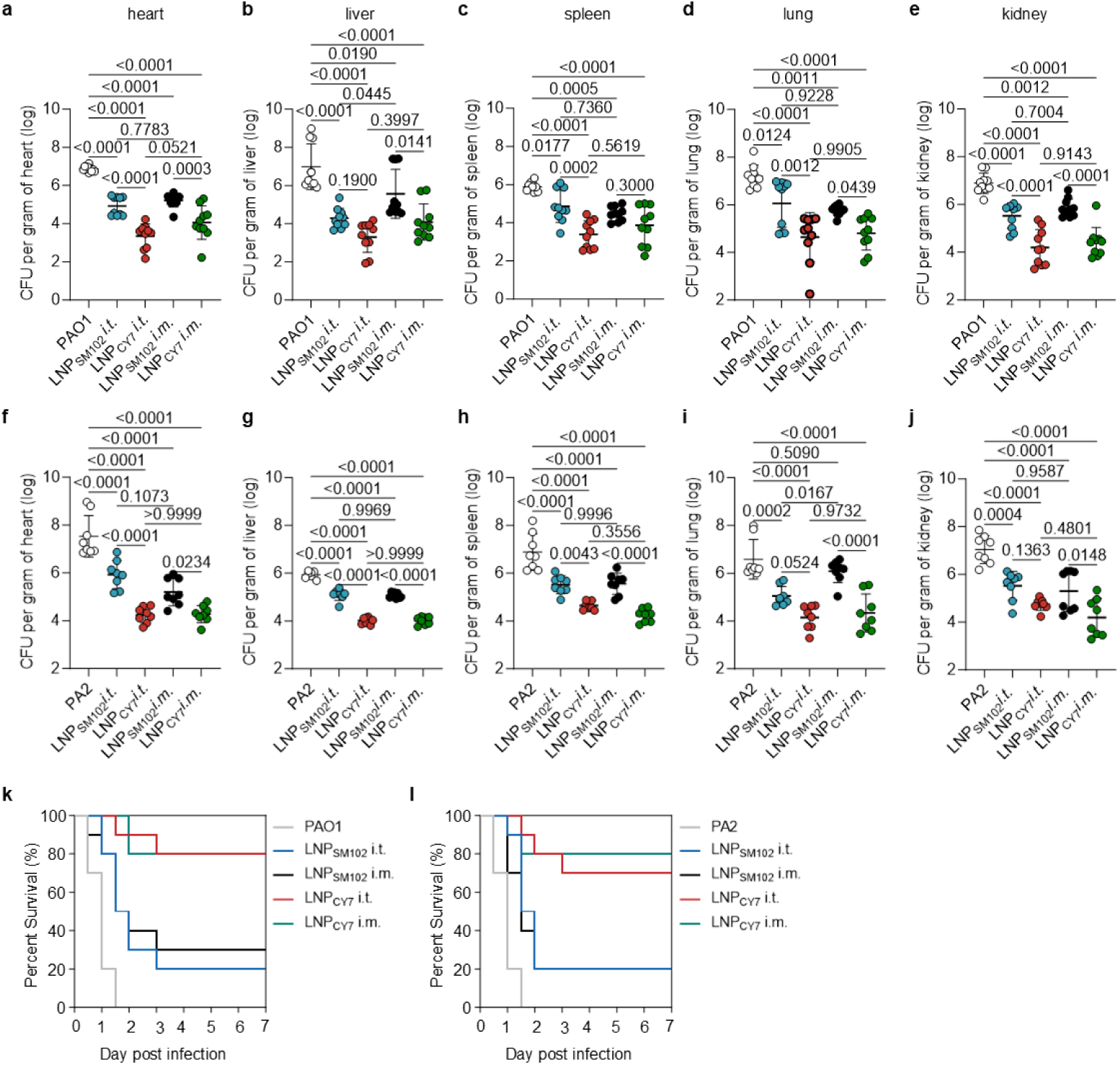
mRNA-LNP_CY7_ vaccination protects against *P. aeruginosa* infections in systemic infection model. **a-e,** Bacteria colony-forming unit (CFU) enumeration per gram of heart (a), liver (b), spleen (c), lung (d) and kidney (e) after challenge with PAO1 in systemic infection model (n=10). **f-j,** Bacteria colony-forming unit (CFU) enumeration per gram of heart (f), liver (g), spleen (h), lung (i) and kidney (j) after challenge with clinical strain PA2 in systemic infection model (n=8). **k,l,** Survival rates of mice (n = 10) immunized with mRNA-LNP vaccines following challenge with either PAO1 strain (k) or the clinical PA2 strain (l) in a systemic infection model. Data are presented as means ± s.d. Statistical significance was determined using one-way ANOVA with Tukey’s multiple comparisons test in GraphPad Prism 10.

To further assess the safety of intratracheal mRNA-LNP administration, we conducted comprehensive evaluations. Histopathological analysis of the lungs, heart, liver, spleen, and kidneys showed no evidence of tissue injury, inflammation, or hemorrhage post-immunization (Supplementary Fig. S18). Serum biochemical markers of hepatic (ALT, AST) and renal (BUN, creatinine) function remained within physiological ranges (Supplementary Fig. S19), and systemic cytokine profiling revealed no abnormal elevations in IL-6, TNF-α, or IL-1β 24 hours post-immunization (Supplementary Fig. S20), collectively supporting a favorable safety profile. Taken together, these results demonstrate that mRNA-LNP_CY7_ vaccination elicits safe and potent antigen-specific adaptive immune responses capable of protecting against both pulmonary and systemic *P. aeruginosa* infections.

## Discussion

Bacterial infections continue to pose a major global health challenge, particularly in the context of escalating antibiotic resistance. Despite the success of vaccination in controlling many infectious diseases, effective vaccines remain unavailable for several clinically important bacterial pathogens, including methicillin-resistant *Staphylococcus aureus* (MRSA), *Pseudomonas aeruginosa*, and *Acinetobacter baumannii*. In addition to the difficulty of identifying appropriate bacterial antigens, the limited efficacy of many bacterial vaccines is strongly influenced by adjuvant design and delivery efficiency. In this study, we engineered a novel ionizable cationic lipid CY7, specifically optimized to promote localized and selective high-level mRNA expression in the lung. LNPs formulated with CY7 achieved markedly higher levels of pulmonary expression compared to those formulated with the clinically used lipid SM102, demonstrating superior pulmonary delivery efficiency to the respiratory tract. Notably, LNP_CY7_ also exhibited enhanced adjuvant activity, resulting in higher antigen-specific humoral and cellular responses and improved protective efficacy relative to LNP_SM102_.

While vaccination remains a cornerstone strategy for the prevention and control of infectious diseases, a major limitation is the temporal delay between immunization and the establishment of effective adaptive immunity. This so-called “immunity gap” leaves individuals vulnerable to infection during the early post-vaccination period, particularly in outbreak or exposure-prone settings^10,11^. Bridging this gap with rapid immune protection remains an unmet need in current vaccine strategies. Here, we demonstrate that pulmonary administration of this mRNA-LNP vaccine confers both rapid and durable protection against bacterial lung infection during the critical window, offering an effective strategy to overcome the transient vulnerability associated with conventional immunization.

In this context, pulmonary mRNA-LNP vaccination may be particularly relevant in clinical settings characterized by an acute risk of infection, such as the early period following hospital admission. Hospitalized patients are often exposed to a broad spectrum of bacterial pathogens, and rapid, non-pathogen specific enhancement of pulmonary innate immunity may help reduce susceptibility to early hospital-acquired infections. Moreover, patients with prolonged hospital stays, especially those requiring mechanical ventilation, face a markedly increased risk of opportunistic respiratory infections, including *P. aeruginosa*. In such settings, a vaccine strategy capable of eliciting both immediate innate protection and longer-lasting immunity may offer complementary benefits.

Mechanistically, the lipid nanoparticle (LNP) component functions not only as a carrier for mRNA delivery but also as a potent immunostimulatory adjuvant that activates innate immune pathways^20^. Pulmonary administration of mRNA-LNPs elicited a transient innate immune response in the lung, characterized by brief induction of cytokines and chemokines, including G-CSF, CXCL-1, IL-6, and IFN-β . Following LNP exposure, phagocytes entered a transcriptionally primed state, maintaining enhanced functional activity even after cytokine levels returned to baseline. This primed state equips neutrophils and macrophages with increased phagocytic capacity and heightened responsiveness to bacterial challenge, enabling rapid and effective antibacterial defense during the early “ immunity gap ” prior to the establishment of adaptive immunity.

In addition, mRNA-LNP vaccination promoted the production of IFN-β, consistent with prior reports^54^. Although the role of type I interferons in bacterial infections is highly context-dependent-with studies reporting both protective and detrimental effects^42,48^, our findings here demonstrate a protective role in the setting of *P. aeruginosa* lung infections. Beyond antibacterial effects, the well-established antiviral function of type I interferons suggests broader applicability of this approach. By inducing localized IFN-I responses in the respiratory tract, pulmonary mRNA-LNP vaccination may also confer protection against viral infections during the immunity gap period.

Collectively, our study demonstrates that a novel ionizable lipid enables efficient and localized pulmonary mRNA expression, thereby enhancing both the immunogenicity and protective efficacy of mRNA vaccines. We further identify a previously underappreciated mechanism by which pulmonary delivery of mRNA-LNP vaccines provides immediate immune protection through priming of the phagocytes. In contrast to conventional vaccines that rely primarily on the gradual development of adaptive immunity, this strategy rapidly induces an antigen-independent innate response, characterized by the activation of immune signaling pathways and interferon-stimulated genes in pulmonary innate immune cells. This response primes neutrophils and macrophages toward a functionally enhanced state, promoting efficient pathogen clearance during the early, vulnerable post-vaccination period. Importantly, this innate response is transient and spatially restricted to the lung, with inflammatory mediators returning to baseline levels within a short time frame, suggesting a controlled activation profile that is unlikely to result in excessive or systemic inflammation. In parallel, pulmonary immunization also elicits robust and durable mucosal as well as systemic adaptive immune responses, providing long-term protection against both laboratory and clinically drug-resistant strains of *Pseudomonas aeruginosa*.

Together, these findings highlight a pulmonary mRNA-LNP platform that can coordinate innate and adaptive immune responses to achieve rapid and durable protection. By demonstrating localized modulation of lung immunity, this strategy provides a foundation for further development of vaccines targeting respiratory pathogens, including potential applications beyond bacterial infections.

## Supporting Information

Supporting Information is available from …

## Acknowledgements

This work was financially supported by the National Natural Science Foundation of China (32330058, 82204303, 82125035) and Shanghai Education Commission Major Project (2021-01-07-00-07-E00081).

## Author contributions

X.W. and A.W. contributed equally to this work. X.W., Q.H., Z.Z., L.F. and C.Z. conceived and designed the experiments. X.W. and A.W. performed the majority of the experiments. L.F. synthesized the ionizable lipid. Y.M. studied and analyzed LNP expressions in vivo. Z.Y. designed and carried out the in vitro transcription (IVT) for mRNA synthesis. Y.M., Z.G., and T.D. assisted with antibody measurements and T cell immunity evaluations. Y.S., K.T. and D.L. contributed to the efficacy studies. G.L. assisted with safety evaluations. X.W. and A.W. analyzed the data and drafted the manuscript. All authors read and revised the manuscript.

## Conflicts of interest

The authors declare no conflict of interest.

## Data availability statement

All data supporting the findings of this study are available within the Article and its Supplementary Information. Source data are provided with the manuscript in the associated source data file. All other data are available from the corresponding author upon reasonable request. The single-cell RNA sequencing (scRNA-seq) data will be deposited in a public repository.

## Code availability statement

All code used in this study has been provided as a supplementary file and submitted with the manuscript.

## Methods

### Materials

SM-102, DSPC, cholesterol, PEG2000-DMG were purchased from A.V.T. Pharmaceutical Co., Ltd. (Shanghai, China). Luc mRNA was purchased from Catug Biotechnology Co., Ltd. (Su Zhou, China). BCA Protein Assay Kit (Cat#71285) were from Sigma (St. Louis, MO). Type IV collagenase were from maokangbio Co., Ltd. (Shanghai, China). TMB (3,3’,5,5’-Tetramethylbenzidine) chromogen solution, DAPI were acquired from Beyotime Biotechnology (Nantong, China). Gradient precast polyacrylamide gels (4–20%) (Cat# 456-1093) and protein dual color standards (Cat# 1610374) were from BIO-RAD (Hercules, CA). Fetal bovine serum (FBS), DMEM, 1640 RPMI, penicillin-streptomycin, D-Luciferin, cell lysis buffer and protease inhibitor were purchased from Dalian Meilun Biotechnology Co., Ltd. (Dalian, China). Cell culture plates and centrifuge tubes were obtained from NEST Biotechnology Co., Ltd. (Wuxi, China). LB (Luria Broth) was purchased from Beijing Anboxing Biotech Co., Ltd. (Beijing, China). Quant-iT™ RiboGreen™ RNA Assay Kit and anti-Mo IFNAR1 antibody (MAR1-5A3) were purchased from Thermo Scientific Co., Ltd. (Rockford, IL, USA). GoldBand DL5,000 DNA Marker, Goldview Nucleic Acid Gel Stain (10,000×) and SDS-PAGE sample loading buffer (5×) were purchased from Yesen (Shanghai, China). Mouse IFN-β ELISA Kit was obtained from Elabscience Biotechnology CO., Ltd. (Wuhan, China). Brilliant Violet 421™ anti-mouse F4/80 Antibody (Cat#123132, clone BM8), APC/Cyanine7 anti-mouse Ly-6C Antibody (Cat#128026, clone HK1.4127628), Brilliant Violet 421™ anti-mouse Ly-6G Antibody (Cat#127628, clone 1A8), Pacific Blue™ anti-mouse/human CD45R/B220 Antibody (Cat#103230, clone RA3-6B2), Alexa Fluor® 647 anti-mouse/human GL7 Antigen Antibody(Cat#144605, clone GL7), FITC anti-mouse CD95 (Fas) Antibody(Cat#152605, clone SA367H8), APC anti-mouse CD3 Antibody (Cat#100236, clone 17A2), Brilliant Violet 421™ anti-mouse CD19 Antibody (Cat#115537, clone 6D5), APC anti-mouse CD11c Antibody (Cat#117310, clone N418), Alexa Fluor® 488 anti-mouse IFN-γ Antibody (Cat#505815, clone XMG1.2), FITC anti-mouse IL-17A Antibody (Cat#506907, clone TC11-18H10.1), FITC anti-mouse TNF-α Antibody (Cat#506303, clone MP6-XT22), Brilliant Violet 421™ anti-mouse CD4 Antibody (Cat#100563, clone RM4-5), APC anti-mouse CD8a Antibody (Cat#100712, clone 53-6.7), Ultra-LEAF™ Purified anti-mouse IFN-β (Cat#508107 clone MIB-5E9.1) mouse IL-1β ELISA, mouse TNF-α ELISA and mouse IL-6 ELISA Kit were purchased from Biolegend. Recombinant Anti-Histone H3(citrulline R2+R8+R17) antibody(ab281584), Anti-Myeloperoxidase antibody(ab300650), and Goat Anti-Mouse IgA alpha chain (HRP) (ab97235) were obtained from Abcam.

### Bacteria, cells and Animals

The standard strain *Pseudomonas aeruginosa* PAO1 and the clinical isolate PA2 were kindly provided by Professor Wuyuan Lu (Fudan University). 293T cells was obtained from the Cell Bank of the Chinese Academy of Sciences (Shanghai, China) and maintained in Dulbecco’s Modified Eagle Medium (Gibco) supplemented with 10% FBS (Gibco), 100 U/mL penicillin, and 100 μg/mL streptomycin at 37 °C under a humidified atmosphere containing 5% CO_2_. Six to eight-week-old male C57BL/6 mice and female BALB/c mice were purchased from GemPharmatech Co., Ltd. (Nanjing, China) and maintained under SPF conditions. Ai9 reporter mice were kindly provided by Professor Yang Hui from the Shanghai Research Center for Brain Science and Brain-Inspired Intelligence. All animal procedures were performed in strict accordance with the Guidelines for the Care and Use of Laboratory Animals of Fudan University, and were approved by the Animal Ethics Committee of Fudan University.

### *In vitro* transcription of PcrV mRNA and OprF-I mRNA

mRNAs encoding PcrV and OprF-I were designed based on published protocols^51^ and synthesized by *in vitro* transcription (IVT). To facilitate protein secretion and detection, a secretory signal peptide was added to the N-terminus of each construct, while a C-terminal 6×His tag was added to PcrV and a C-terminal Flag tag to OprF-I. Coding sequences were cloned into a plasmid vector optimized for mRNA production, containing enhanced 5′ and 3′ untranslated regions (UTRs) and a poly(A) tail signal. The resulting nucleoside-modified mRNAs incorporated N1-methyl-pseudouridine (m1Ψ) UTP, a 100-nucleotide poly(A) tail, and a cap 1 structure using a trinucleotide cap 1 analogue. mRNA was synthesized *in vitro* using the T7 Co-transcription RNA Synthesis Kit (Kactus Biosystems, Shanghai, China), precipitated with LiCl, and purified by cellulose chromatography. The integrity and purity of the synthesized mRNAs were confirmed by agarose gel electrophoresis. A residual double-stranded RNA (dsRNA) contaminants content was 0.09 ± 0.03%, measured by a commercial dsRNA ELISA kit (36717ES48, Yeasen) (Shanghai, China), which is well below the quality standard for residual dsRNA 0.2% (2000 pg/µg) according to the BioNTech and Moderna mRNA manufacturing patents^55^.

293T cells (1×10⁵/well) were seeded in 12-well plates and incubated overnight. For transfection, 1 μL or 2 μL mRNA (1 μg/μL) and 2 μL Lipofectamine 2000 were separately diluted in serum-free DMEM, then combined and incubated at room temperature for 15 minutes to allow complex formation. The transfection mixture was added dropwise to the cells and incubated for 24 hours. After transfection, both the supernatant and cell lysates were collected. Protein samples were separated by 4-20% SDS-PAGE and transferred onto PVDF membranes. Membranes were blocked with 5% non-fat milk in PBST for 1 hour at room temperature, then incubated with HRP-conjugated anti-His antibody or anti-Flag antibody overnight at 4 °C. After washing with 0.1% PBST, signal was visualized using TMB substrate according to the manufacturer’s instructions.

### Preparation and characterization of LNPs

LNPs were prepared by microfluidic mixing of an ethanol phase containing lipids with an aqueous phase containing mRNA, as previously described^56^. Specifically, the ethanol phase comprised ionizable lipid, cholesterol, DSPC, and DMG-PEG at a molar ratio of 50:38.5:10:3. This was mixed with an aqueous phase consisting of 25 mM citrate buffer (pH 5.2) containing mRNA, using a microfluidic device at a flow rate ratio of 1:3 (ethanol:aqueous) and an N/P ratio of 5.67:1. The resulting LNPs were diluted and purified by ultrafiltration using a 100 kDa molecular weight cut-off filter. The hydrodynamic diameter of LNPs was measured using a Zetasizer Nano ZS (Malvern). The mRNA concentration and encapsulation efficiency were determined by RiboGreen assay according to the manufacturer’s protocol. The morphology of the LNPs was visualized using Cryo-EM (JEOL Cryo ARM 300 kV).

### *In vivo* bioluminescence imaging

To assess *in vivo* protein expression, Luc mRNA-loaded LNPs were administered to male C57BL/6 mice or female BALB/c mice via two different routes. For intratracheal nebulization, mice were anesthetized and administered 30 μL of Luc mRNA-LNPs (5 μg mRNA per mouse) using a Micro-Sprayer Aerosolizer. For intramuscular injection, 30 μL of the same formulation was injected into the left thigh muscle. At 6 hours post-administration, mice were injected intraperitoneally with 200 μL of D-luciferin solution (15 mg/mL in PBS). Mice were then anesthetized with isoflurane, and bioluminescence signals were captured 5 minutes after luciferin injection using the VISQUE® InVivo Smart imaging system.

### *In vivo* mRNA delivery and cellular expression analysis

Ai9 reporter mice were administered 30 μL of Cre mRNA-loaded LNPs (2 μg mRNA per mouse) via intratracheal nebulization. At 72 hours post-delivery, mice were euthanized and perfused with 1× PBS, and lungs and livers were harvested. Tissues were fixed overnight in 4% paraformaldehyde (PFA), processed in sucrose, and embedded in OCT compound. Frozen sections (15 μm thickness) were prepared, mounted using anti-fade mounting medium containing DAPI, and imaged with confocal microscopy to visualize Cre-mediated tdTomato expression.

In parallel experiments, lungs and mediastinal lymph nodes were collected for cellular-level mRNA expression analysis. Mice were euthanized and perfused with ice-cold Hank’s Balanced Salt Solution (HBSS). Lungs were dissected, finely minced with sterile scissors, and enzymatically digested in 1 mg/mL Type IV collagenase and 10 μg/mL DNase I at 37 °C for 30 minutes. The resulting cell suspensions were passed through 70 μm cell strainers, centrifuged at 400 × g for 10 minutes, and subjected to red blood cell (RBC) lysis. After incubation on ice for 5 minutes, the lysis reaction was quenched with 10 volumes of cold 1× PBS, followed by another centrifugation step. Cell pellets were resuspended in 1 mL of 1% BSA in PBS, stained with fluorochrome-conjugated antibodies for 30 minutes at 4 °C, washed with PBS, and analyzed using a flow cytometer.

To assess cell-type specific uptake of mRNA-LNP, 30 μL of Cy5 labeled mRNA-LNP were administered via intratracheal nebulization (2 μg mRNA per mouse). At 6 and 24 hours post-delivery, mice were euthanized and single cells were harvested as above. The cellular uptake was further analyzed using a flow cytometer.

### IL-6 quantification in lymph nodes

Female BALB/c mice were immunized with 2 μg of PcrV mRNA-loaded LNPs (LNP_CY7_ or LNP_SM102_) via intratracheal nebulization or intramuscular injection. At designated time points post-immunization, mediastinal lymph nodes (intratracheal group) or iliac lymph nodes (intramuscular group) were harvested and mechanically homogenized at 65 Hz for 1 minute for cytokine analysis.

IL-6 levels in the lymph node homogenates were measured using a commercial ELISA kit according to the manufacturer’s instructions. Briefly, 96-well plates were coated with 50 μL of capture antibody diluted in ELISA coating buffer and incubated overnight at 4 °C. After four washes with 0.05% PBST, plates were blocked with 1% BSA for 2 hours at room temperature. Lyophilized standards were reconstituted at 500 pg/mL and subjected to 2-fold serial dilutions. Samples and standards (100 μL each) diluted in blocking buffer were added to the wells and incubated for 2 hours at room temperature with shaking. After washing, 100 μL of biotinylated detection antibody was added and incubated for 1 hour, followed by 100 μL of avidin-HRP. After three additional washes, TMB substrate was added for color development, and the reaction was stopped using 2 N H₂SO₄. Absorbance was measured at 450 nm using a microplate reader.

### Germinal center formation

Female BALB/c mice were immunized with PcrV mRNA-loaded LNP_CY7_ or LNP_SM102_ (2 μg mRNA per dose) via intratracheal nebulization or intramuscular injection on days 0, 7, and 21. Seven days after the final immunization, mediastinal lymph nodes (intratracheal group) or iliac lymph nodes (intramuscular group) were harvested and processed into single-cell suspensions. Lymph nodes were mechanically dissociated, filtered through a 70 μm cell strainer, and centrifuged at 500 × g for 5 minutes. Cell pellets were resuspended in 1% BSA in PBS and stained at 4 °C for 30 minutes with a fluorescent antibody cocktail containing: FITC-conjugated anti-mouse CD95 (Fas), Alexa Fluor® 647-conjugated anti-mouse/human GL7 and Pacific Blue™-conjugated anti-mouse/human CD45R/B220. After staining, cells were washed to remove unbound antibodies and analyzed using a flow cytometer (Agilent) to evaluate germinal center B cell populations.

### Determination of antigen-specific antibody titers

Female BALB/c mice were immunized via intratracheal nebulization or intramuscular injection with LNP_CY7_ or LNP_SM102_ formulations containing 1 μg each of PcrV mRNA and OprF-I mRNA per dose on days 0, 7, and 21. Serum samples were collected on days 7, 14, and 28 for quantification of anti-PcrV and anti-OprF-I IgG antibody titers. On day 35, mice were euthanized, and bronchoalveolar lavage fluid (BALF) was collected to measure secretory IgA levels. For antibody detection, 96-well ELISA plates were coated overnight at 4 °C with 100 μL/well of recombinant PcrV or OprI protein (expressed in *E. coli*). Plates were washed three times with 0.1% PBST, then blocked with 1% BSA in PBS for 2 hours at room temperature. After additional washes, serially diluted serum or BAL samples were added to the wells and incubated for 2 hours at room temperature with gentle shaking. Plates were then washed and incubated with HRP-conjugated goat anti-mouse IgG (for serum) or IgA (for BALF) antibodies (1:5000 dilution) for 2 hours. After four washes, 100 μL of TMB substrate was added to each well for color development, and the reaction was stopped after 10 minutes with 100 μL of 2 N H₂SO₄. Absorbance was measured at 450 nm using a microplate reader.

### ELISpot assay

Following the vaccination schedule described above, spleens were harvested on day 28 and processed into single-cell suspensions by gentle mechanical dissociation in digestion buffer containing 1 mg/mL Type IV collagenase and DNase. The cell suspension was filtered through a 70 μm cell strainer, centrifuged at 400 × g for 5 minutes, and resuspended in RBC lysis buffer. After 5 minutes of incubation on ice, the reaction was quenched with 10 volumes of cold 1× PBS, followed by centrifugation at 400 × g for 5 minutes at 4°C. The resulting cell pellets were resuspended in RPMI 1640 medium and plated onto pre-activated, pre-coated 96-well ELISpot plates at a density of 3.6 × 10⁵ cells per well. For antigen-specific stimulation, experimental wells were treated with 0.2 μg of each recombinant protein and incubated for 24 hours at 37°C with 5% CO₂. Control wells received either PMA (positive control) or culture medium alone (negative control). After incubation, cells were lysed with cold deionized water and plates were washed five times with PBST buffer. The detection procedure consisted of incubation with 100 μL of 1× biotinylated primary antibody at 37°C for 1 hour, followed by washing and incubation with streptavidin-HRP conjugate at 37°C for 1 hour. Plates were washed five times again before developing spots with TMB substrate for 30 minutes in the dark. The reaction was stopped by washing with deionized water, and plates were air-dried. Antigen-specific spot-forming units were enumerated using an automated ELISpot reader.

### Mouse Immunization and evaluation of rapid protection

Female BALB/c mice were immunized via intratracheal nebulization or intramuscular injection with PcrV mRNA-loaded LNP_CY7_ or LNP_SM102_ formulations (2 μg mRNA per mouse). Three days post-immunization, mice were anesthetized and challenged intratracheally with either *Pseudomonas aeruginosa* PAO1 (2 × 10^6^ CFU/mouse) or PA2 (1 × 10^6^ CFU/mouse). Twenty-four hours after bacterial challenge, lungs were harvested and homogenized in 1 mL sterile PBS using bead-beating (60 Hz, 3 min). Homogenates were serially diluted in PBS, and 30 μL aliquots of each dilution were plated on LB agar plates. After overnight incubation at 37°C, bacterial colonies were counted and expressed as colony-forming units per gram /lung.

In a parallel experiment, mice immunized under the same conditions were challenged with higher bacterial doses (1 × 10^7^ CFU PAO1 or 5 × 10^6^ CFU PA2 per mouse), and survival was monitored over time.

For histopathological analysis, lung tissues were collected 24 hours post-infection, fixed in 4% PFA for 24 hours, processed through graded ethanol series, embedded in paraffin, and sectioned at 5 μm thickness. Sections were stained with hematoxylin and eosin (H&E) to evaluate pulmonary inflammation, immune cell infiltration, and tissue damage.

To assess the protection conferred by mRNA-LNP formulations, additional control groups were included. Mice received intratracheal nebulization of the following preparations (all doses normalized to contain an equivalent of 2 μg active component): empty LNP (containing 30 μg ionizable cationic lipid, equivalent to lipid content in 2 μg mRNA-LNP), 2 μg free PcrV mRNA, 2 μg PcrV mRNA + empty LNP, and 2 μg PcrV mRNA-LNP. On day 3, all groups were challenged intratracheally with 2 × 10^6^ CFU PAO1, and bacterial burden was quantified as described above.

To determine the functional contribution of specific innate cell types to early protection, neutrophils were depleted by tail vein injection of 150 μg Ultra-LEAF™ anti-Ly6G antibody^35,36^, whereas macrophages were depleted by intraperitoneal administration of 200 μL clodronate liposomes (5 mg/mL)^37,38^. Flow cytometric analysis confirmed efficient depletion of the targeted myeloid subsets in the lungs 24 h after treatment. BALB/c mice were vaccinated intratracheally with mRNA-LNP_CY7_ and subjected to cell depletion 48 h later. At 24 h post-depletion, mice were challenged with 2 × 10^6^ CFU PAO1, and lung bacterial burdens were quantified 24 h after infection.

To investigate the duration of innate immune responses, bacterial enumeration in PAO1-infected lungs was assessed 24 hours after pulmonary infection on days 1, 3, 5, 7, and 10 post-vaccination (with a booster administered on day 7).

To evaluate the breadth of protection, an additional cohort of immunized mice was challenged intratracheally with *Acinetobacter baumannii* (2 × 10^6^ CFU/mouse) on day 3 post-vaccination. Lung bacterial loads were assessed 24 hours post-challenge.

To further investigate the protective mechanism, mice were assigned to three groups: (1) naive (unvaccinated), (2) vaccinated (PcrV mRNA-LNP), and (3) antibody-blocked (vaccinated with anti IFN-β antibody). In the antibody-blocked group, anti-IFN-β monoclonal antibody (40 μg) was co-administered intratracheally with PcrV mRNA-LNP during immunization. Three days post-vaccination, all mice were challenged intratracheally with PAO1 (2 × 10^6^ CFU), and lungs were harvested 24 hours later for bacterial enumeration.

### Single-cell RNA sequencing (scRNA-seq)

Female BALB/c mice were anesthetized and intratracheally nebulized with PcrV mRNA-loaded LNP_CY7_ (2 μg mRNA per mouse) using a microsprayer. Three days post-immunization, mice were euthanized, and lungs were collected along with control lungs. The tissues were immediately transferred into tissue storage buffer on ice, then washed in 10 mL of chilled 1× Dulbecco’s Phosphate-Buffered Saline (DPBS) to remove residual buffer. Tissues were finely minced on ice and enzymatically digested in a solution containing Collagenase (1 mg/mL), Dispase (1 mg/mL), and DNase I (10 μg/mL) at 37 °C for 30 minutes with gentle agitation. The resulting cell suspensions were filtered through 40 μm cell strainers and subjected to red blood cell (RBC) lysis^57^. After lysis, cells were washed with 1× DPBS supplemented with 0.4% fetal bovine serum (FBS). Cell viability was assessed using 0.4% acridine orange/propidium iodide (AO/PI) staining and analyzed on the Countstar Rigel S2 system. Viable single-cell suspensions were submitted to Majorbio Biopharm Technology Co., Ltd. (Shanghai, China) for single-cell RNA sequencing.

Raw sequencing data generated on the Illumina platform were converted to FASTQ files and processed using the Cell Ranger software (v7.1.0) with default parameters. Reads were aligned to the mouse reference genome using the STAR algorithm. Unique molecular identifiers (UMIs) were counted, and cell barcodes not corresponding to real cells were filtered out, resulting in a filtered gene expression matrix. The gene-barcode matrices were imported into Seurat (v4.4.0) for downstream analysis^58^. Cells were filtered based on the following quality control criteria: (1) fewer than 200 or more than 6,000 detected genes, or (2) mitochondrial gene content exceeding 5%. Raw UMI counts were log-normalized, and 2,000 highly variable genes were selected. These were scaled using the ScaleData() function^54^. Subsequently, dimensionality reduction was performed using principal component analysis (PCA), followed by graph-based clustering and cell type annotation following standard Seurat workflows.

For cell-type annotation, canonical marker genes from the literature were used to define major lung cell populations. A dot plot was generated to visualize the expression of these marker genes across clusters. In the plot, dot size represents the proportion of cells in each cluster expressing a given gene, while color reflects the min-max normalized average expression. Based on these patterns, clusters were manually annotated to specific cell types.

Differential gene expression analysis between neutrophils from different groups was performed using DESeq2. Genes with an adjusted p-value < 0.05 and a log_2_ fold change > 0.5 were considered significantly differentially expressed. Volcano plots were generated to visualize the results. To further examine ISG expression in neutrophils, we calculated ISG signature scores using the AddModuleScore function in Seurat, based on a curated list of canonical ISGs. Cells in the top 10% of ISG scores were defined as ISG^hi,^ and the rest as non-ISG^hi^. DEGs between ISG^hi^ and non-ISG^hi^ neutrophils were identified using FindMarkers with a log_2_ fold change threshold of 0.25 and minimum expression in 10% of cells. The top DEGs were visualized in a heatmap using DoHeatmap.

### NETs formation

In a parallel experiment under the same immunization conditions, mice were intratracheally challenged with *Pseudomonas aeruginosa* PAO1 (2 × 10^6^ CFU) on day 3 post-vaccination. Twenty-four hours after infection, mice were euthanized and lungs were harvested. Lung tissues were fixed in 4% paraformaldehyde, dehydrated through a graded sucrose series (20-30%), and embedded in OCT compound. Cryosections of 15 μm thickness were prepared and subjected to immunofluorescence staining. Sections were blocked with 5% bovine serum albumin (BSA) for 1 hour at room temperature, then incubated overnight at 4°C with primary antibodies against citrullinated histone H3 (CitH3, 1:200) and myeloperoxidase (MPO, 1:200). After washing, sections were incubated with fluorophore-conjugated secondary antibodies for 1 hour at room temperature. Nuclei were counterstained with DAPI (1 μg/mL), and slides were mounted with anti-fade mounting medium. NET formation was visualized using a laser scanning confocal microscope (Leica MICA).

### Mouse immunization and evaluation of long-term adaptive protection

Female BALB/c mice were immunized via intratracheal nebulization or intramuscular injection with a mixture of PcrV mRNA-LNP and OprF-I mRNA-LNP. Each dose contained 1 μg of PcrV mRNA and 1 μg of OprF-I mRNA and was administered on days 0, 7, and 21 following a prime-boost schedule. On day 35, mice were intratracheally challenged with *Pseudomonas aeruginosa* strains PAO1 (2 × 10^6^ CFU/mouse) or PA2 (1 × 10^6^ CFU/mouse). Twenty-four hours post-challenge, bacterial loads in lung tissues were quantified by colony-forming unit (CFU) enumeration as previously described. For survival analysis, mice were challenged intratracheally with higher doses of PAO1 (1 × 10^7^ CFU/mouse) or PA2 (5 × 10^6^ CFU/mouse), and survival was monitored daily for 7 days post-infection.

To assess systemic protection, vaccinated mice were intravenously challenged with PAO1(5 × 10^7^ CFU/mouse) or PA2 (1 × 10^7^ CFU/mouse) in 100 μL sterile PBS. Twenty-four hours after intravenous challenge, major organs (liver, lung, heart, spleen, and kidneys) were harvested, weighed, and homogenized in sterile PBS at a ratio of 1 mL per 100 mg tissue. Tissue homogenates were serially diluted and plated on LB agar plates for bacterial CFU quantification (CFU per gram of tissue). In parallel, a survival study was conducted by monitoring mice for 7 days following intravenous bacterial challenge.

### Safety evaluation

For dose-escalation study, female BALB/c mice were intratracheally administered with mRNA-LNP containing 1, 2 or 5 μg mRNA on days 0, 7 and 21. Control mice received PBS only. Body weight was recorded every other day. Lung tissues were collected 24 hours post-administration and subjected to HE staining.

Following the vaccination protocol described above, serum samples were collected 24 hours after each immunization for comprehensive safety assessment. Hepatic and renal function biomarkers-including alanine aminotransferase (ALT), aspartate aminotransferase (AST), creatinine (CRE-2), and urea nitrogen (UN)-were measured using an automated biochemistry analyzer (ADVIA Chemistry XPT) according to the manufacturer’s instructions.

Simultaneously, serum cytokine levels, including interleukin-6 (IL-6), tumor necrosis factor-alpha (TNF-α), and interleukin-1 beta (IL-1β), were quantified using commercial ELISA kits (BioLegend) following standardized protocols. For histopathological analysis, tissues were collected, fixed, and processed for hematoxylin and eosin (H&E) staining as previously described.

### Statistical analysis

Prism (Graphpad Software) was used to perform statistical analysis. One-way or two-way Anova with Tukey’s multiple comparisons test were adopted to determine differences. Data are presented as mean ± standard deviation (s.d.). In all analysis, p < 0.05 was considered statistically significant (n.s.: non-significance, *p < 0.05, **p <0.01, ***p < 0.001).

## Notes

### Competing Interest Statement

The authors have declared no competing interest.

