## Supplementary Information for "Pulmonary mRNA-LNP Vaccines for Rapid and Durable Protection Against Bacterial Infection"

### 1. Synthesis of ionizable lipids.

*(1) Synthesis of **MA** series ionizable lipids:*

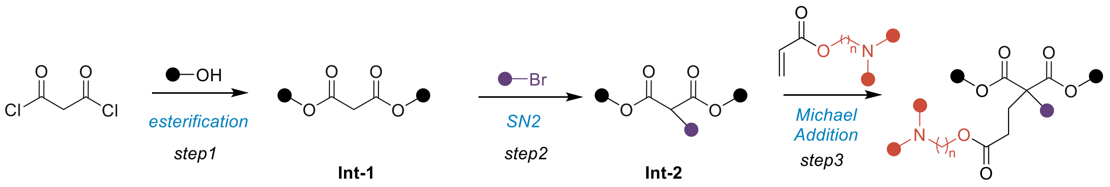


Scheme 1: synthesis of MA series ionizable lipids:

Step 1: The specific alcohol (11 mmol, 2.2 equiv.) was added dropwise to a stirred suspension of malonyl chloride (5 mmol, 1.0 equiv.) in dry DCM (50 mL) at 0 °C under a nitrogen atmosphere. Then $Et_3N$ (15 mmol, 3.0 equiv.) was added, and the mixture was stirred for 8 hours. After removing the organic solvent under vacuum, the crude product was purified by column chromatography on silica gel (PE/EtOAc = 95:5) to afford **Int-1** (65 - 78% yield) as colorless oil.

Step 2: To a solution of **Int-1** (3 mmol, 1.0 equiv.) in anhydrous THF (20 mL) at room temperature was added NaH (3 mmol, 1.1 equiv.), then the reaction mixture was stirred at room temperature for 30 minutes. Then different alkyl bromides (3.3 mmol, 1.1 equiv.) were added and the reaction was moved to 60 °C. 16 hours later, the reaction was concentrated in vacuo and the crude product was purified by column chromatography on silica gel (PE/EtOAc = 96:4) to afford **Int-2** as colorless oil (40% - 73% yield).

Step 3: To a solution of **Int-2** (0.2 mmol, 1.0 equiv.) in anhydrous toluene (2 mL) at room temperature was added NaH (0.06 mmol, 0.3 equiv.), then the reaction mixture was stirred at room temperature for 10 minutes. Then different amino-substituted acrylates (0.3 mmol, 1.5 equiv.) were added and the reaction was moved to 40 °C. 12 hours later, the reaction was concentrated in vacuo and the crude product was purified by column chromatography on silica gel (DCM/MeOH = 94:6) to afford desired **MA** products as colorless oil (50% - 84% yield).

*(2) Synthesis of **CY** series ionizable lipids:*

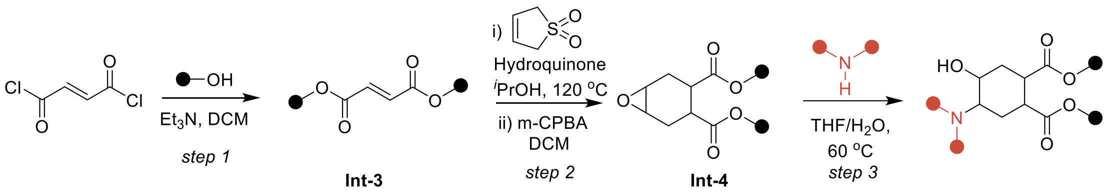


Scheme 2: synthesis of CY series ionizable lipids:

Step 1: The alcohol (11 mmol, 2.2 equiv.) was added dropwise to a stirred suspension of fumaryl chloride (5 mmol, 1.0 equiv.) in dry DCM (50 mL) at 0 °C under a nitrogen atmosphere. Then $Et_3N$ (15 mmol, 3.0 equiv.) was added, and the mixture was stirred for 8 hours. After removing the organic solvent under vacuum, the crude product was purified by column chromatography on silica gel (PE/EtOAc = 95:5) to afford **Int-3** (78% yield) as colorless oil.

Step 2: To a solution of **Int-3** (3 mmol, 1.0 equiv.) and 3-sulfolene (4.5 mmol, 1.5 equiv.) in anhydrous $^{i}$PrOH (20 mL) at room temperature was added hydroquinone (0.3 mmol, 0.1 equiv.), then the reaction mixture was stirred at 120 °C for 24 hours. When **Int-3** was completely consumed, the reaction was concentrated in vacuo and the crude product was purified by column chromatography on silica gel (PE/EtOAc = 95:5) to afford the ester as colorless oil. Then the ester was dissolved in DCM (20 mL) followed by addition of *m*-CPBA (1.2 equiv.). After stirring at room temperature for 12 hours, the organic solvent was concentrated in vacuo and the crude product was purified by column chromatography on silica gel (PE/EtOAc = 92:8) to afford **Int-4** as colorless oil.

Step 3: To a solution of **Int-4** (0.2 mmol, 1.0 equiv.) in THF/$H_2O$ (2 mL, 9:1 v/v) at room temperature was added amine (6 mmol, 30 equiv.), then the reaction mixture was stirred at 60 °C for 24 hours. When **Int-4** was completely consumed, the reaction was concentrated in vacuo and the crude product was purified by column chromatography on silica gel (DCM/MeOH = 95:5) to afford desired **CY** products as colorless oil (56 -

82% yield).

*(3) Synthesis of bis(2-octyldodecyl) 4-(4-(dimethylamino)piperidin-1-yl)-5-hydroxycyclohexane-1,2-dicarboxylate (CY7)*

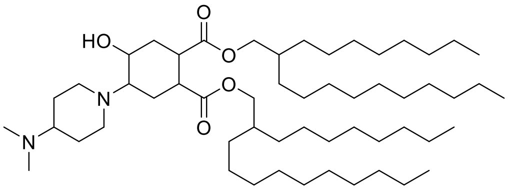


**$^1$H NMR** (600 MHz, Chloroform-*d*) δ 4.08 – 3.94 (m, 4H), 3.75 (s, 1H), 3.42 (td, $J$ = 10.7, 4.2 Hz, 1H), 3.33 – 3.25 (m, 2H), 2.79 (d, $J$ = 11.3 Hz, 1H), 2.68 (d, $J$ = 11.4 Hz, 1H), 2.63 – 2.57 (m, 1H), 2.56 – 2.51 (m, 1H), 2.27 (s, 6H), 2.22 (d, $J$ = 12.7 Hz, 2H), 2.09 – 2.02 (m, 1H), 1.82 (t, $J$ = 12.3 Hz, 2H), 1.72 – 1.46 (m, 8H), 1.33 – 1.21 (m, 63H), 0.88 (t, $J$ = 6.9 Hz, 12H). **HRMS** (ESI) m/z $(M+H)^+$ calculated for $C_{55}H_{107}N_2O_5$: 875.8180, observed: 875.8246.

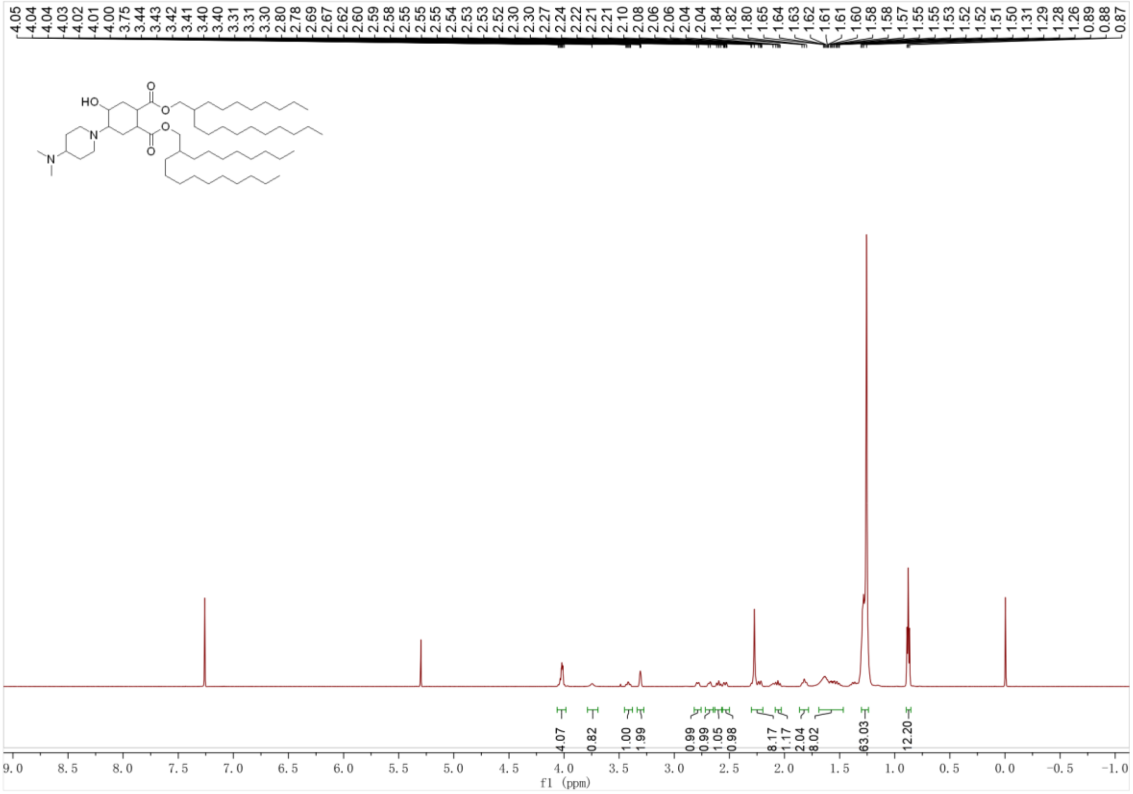


$^1$H-NMR spectrum of **CY7**

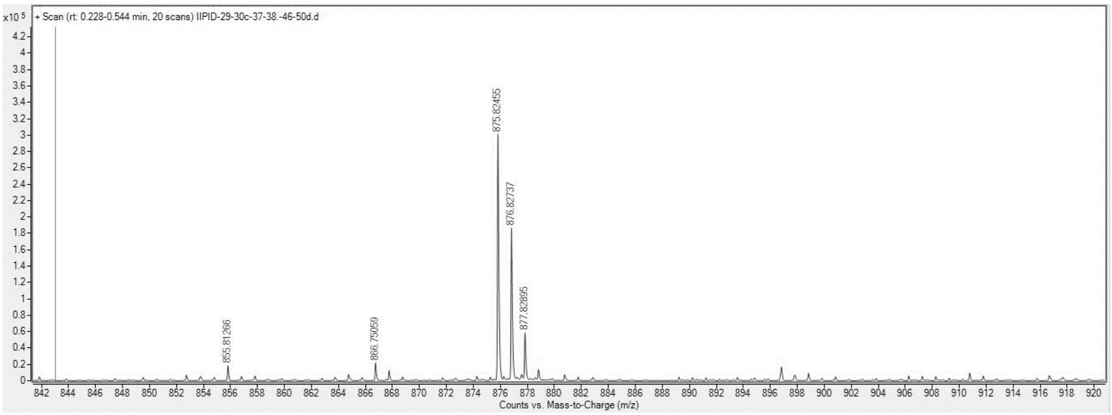


HRMS spectrum of **CY7**

### 2. Supplementary figures.

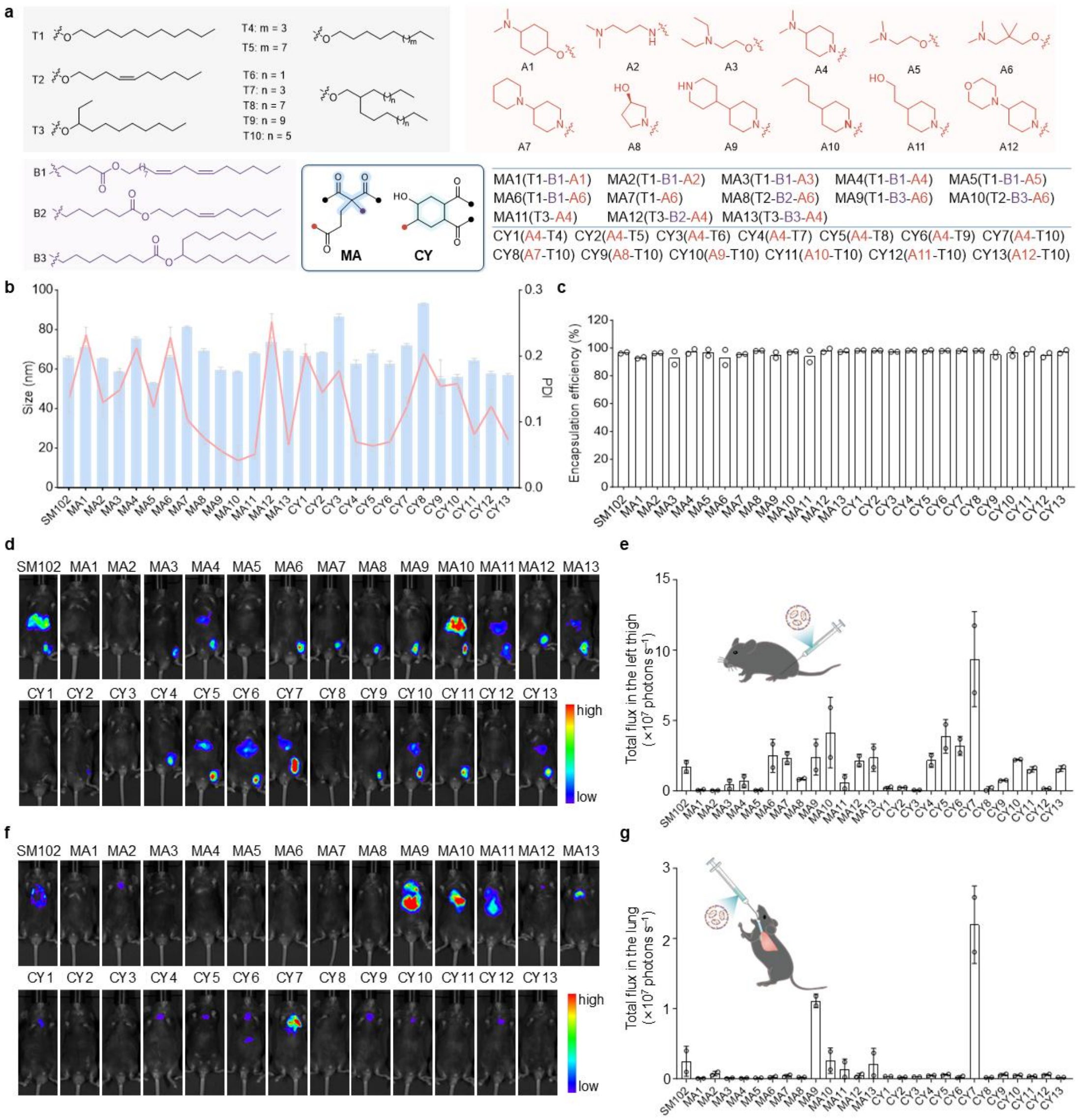


Supplementary Figure S1. Design and evaluation of ionizable lipids for high localized mRNA expression. a) Schematic illustration of two lipid series (MA1-13 and CY1-13) synthesized using malonate and cyclohexane as structural cores. b) Z-average diameter and PDI of all LNP formulations measured by Dynamic Light Scattering (DLS) (n=3). c) mRNA encapsulation efficiency determined by RiboGreen assay (n=2). d) Representative whole-body bioluminescence imaging at 6 hours post-treatment with Luc mRNA-LNPs following intramuscular injection (i.m.) e) Quantification of total flux at the injection site after intramuscular injection (n=2). f) Representative whole-body bioluminescence imaging at 6 hours post-treatment with Luc mRNA-LNPs following intratracheal nebulization (i.t.). g) Quantification of total flux in the lung after intratracheal nebulization (n=2). Data are presented as mean ± s.d.

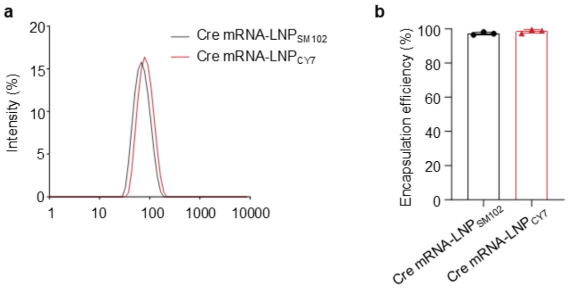


Supplementary Figure S2. Preparation and characterization of Cre mRNA-LNPs. a) Z-average diameter measured by dynamic light scattering (DLS). b) Encapsulation efficiency of Cre mRNA-LNPs determined by RiboGreen assay (n = 3). Data are presented as mean ± s.d.

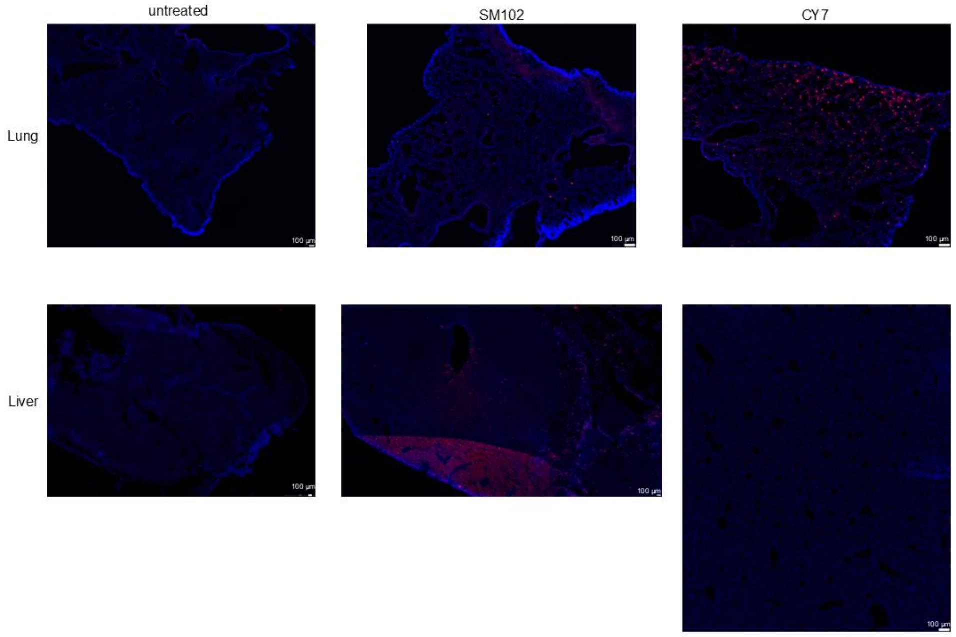


Supplementary Figure S3. Raw fluorescence images of lung and liver tissue sections from Ai9 reporter mice, 3 days after intratracheal administration of Cre mRNA-LNPs formulated with CY7 or SM102, acquired by panoramic scanning on the Mica Microhub (Leica Microsystems (Shanghai) Trading Co., Ltd.). Scale bar = 100 μm.

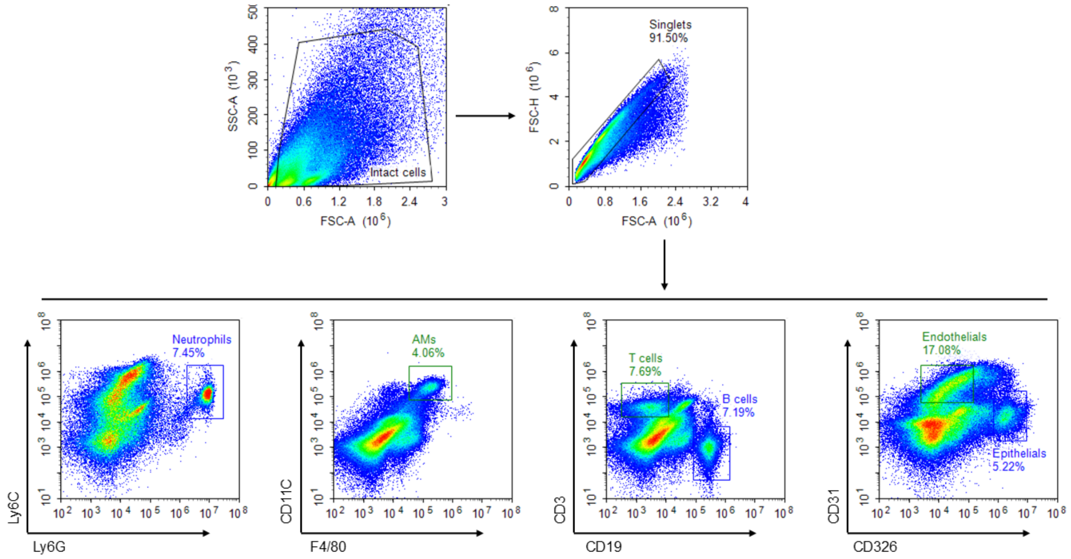


Supplementary Figure S4. Flow cytometry gating strategy for lung cell analyses. Lung single-cell suspensions were first gated to exclude debris, followed by singlet discrimination to ensure analysis of intact cells. Immune and non-immune cell subsets were then identified based on established marker combinations: neutrophils (Ly6G$^+$ Ly6C$^{int}$), alveolar macrophages (F4/80$^+$ CD11c$^+$), B cells (CD19$^+$), T cells (CD3$^+$), endothelial cells (CD31$^+$ CD326$^-$), and epithelial cells (CD326$^+$ CD31$^-$). The same gating pipeline was applied consistently across all experiments and figures.

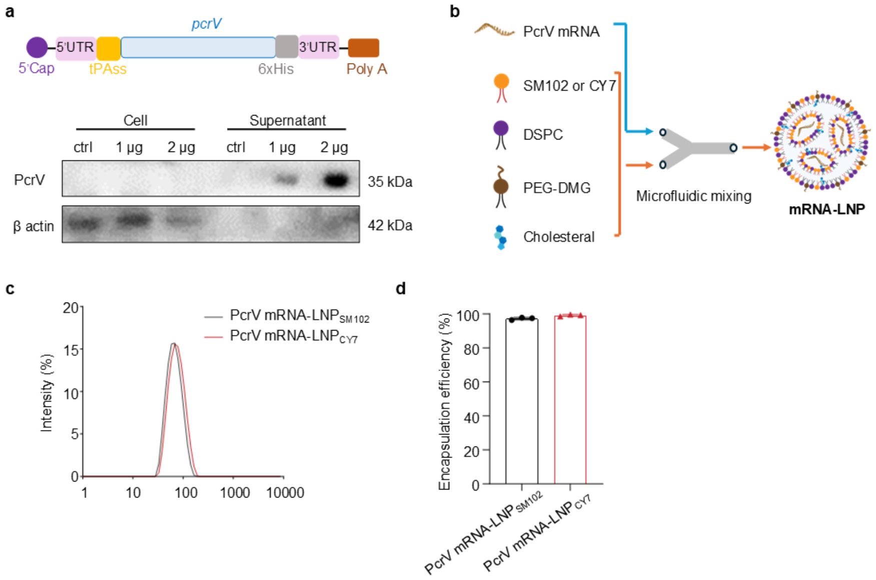


Supplementary Figure S5. PcrV mRNA-LNP fabrication and characterization. a) Schematic representation of the PcrV mRNA construct, consisting of a 5′cap, 5′untranslated region (5′UTR), signal peptide, coding sequence of PcrV, 3′UTR, and a poly(A) tail. Western blot analysis of PcrV protein expression and secretion 24 hours after transfection of HEK-293T cells with 1 μg or 2 μg PcrV mRNA. The results indicate that the protein was predominantly expressed and secreted into the culture supernatant. b) Schematic representing the preparation of PcrV mRNA-LNP. c) Particle size measured by dynamic light scattering (DLS). d) Encapsulation efficiency of PcrV mRNA-LNPs determined by RiboGreen assay (n = 3).

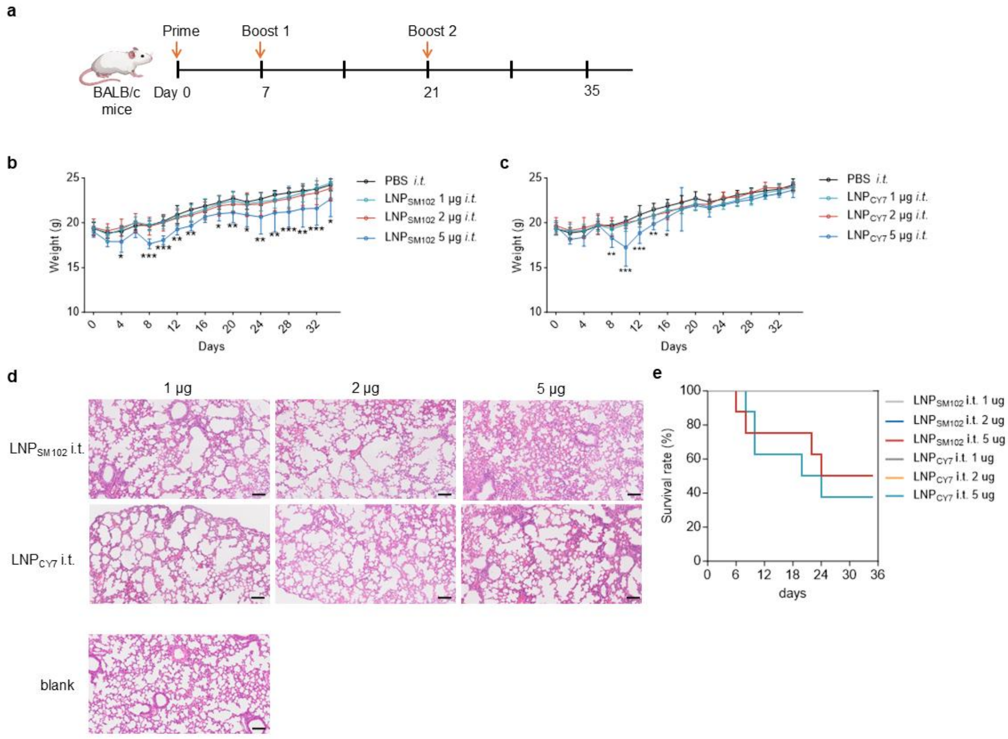


Supplementary Figure S6. Dose-escalation study of pulmonary mRNA-LNP vaccination. a) Timeline of mouse immunization. b and c) Body weight monitoring following intratracheal administration of mRNA-LNPs containing 1 μg, 2 μg, or 5 μg of mRNA. Control mice received PBS only. High-dose (5 μg) administration led to significant weight loss after each administration. d) Representative lung tissue sections HE staining 24 hours post-administration, showing pronounced acute inflammation and marked infiltration of inflammatory cells in 5 μg-treated mice, whereas 1 μg, 2 μg, and control mice exhibited no obvious tissue damage. Scale bar=100 μm. e) Survival curve of mice following mRNA-LNP administration. Data are presented as mean ± s.d. (n = 8). Statistical significance was determined using two-way ANOVA with Tukey's multiple comparisons test in GraphPad Prism 10. * indicates $p < 0.05$; ** indicates $p < 0.01$; *** indicates $p < 0.001$.

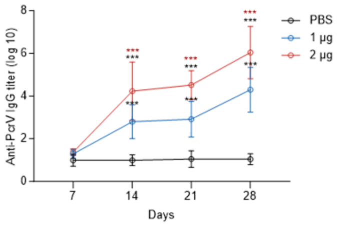


Supplementary Figure S7. PcrV-specific IgG antibody titers in the serum measured after PcrV mRNA-LNP$_{CY7}$ vaccination, demonstrating higher titers in mice receiving 2 μg mRNA compared with 1 μg. Data are presented as mean ± s.d. (n=8). Statistical analyses were performed using two-way ANOVA in Prism 10. Black asterisks indicate comparisons between each dose group and the PBS control, and red asterisks indicate comparisons between the 2-μg and 1-μg groups. *** indicates p < 0.001.

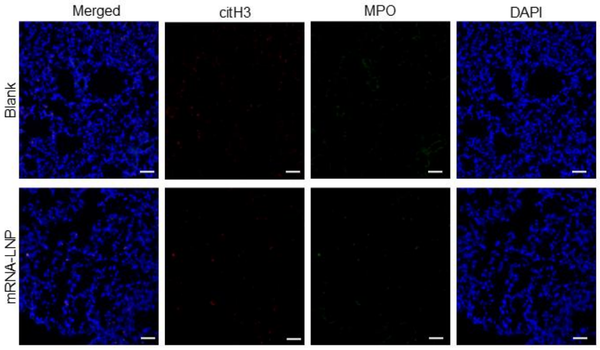


Supplementary Figure S8. Representative immunofluorescence images of the lungs from vaccinated mice 72 h post-vaccination, showing no detectable NET formation. Citrullinated histone H3 (Cit-H3) is shown in red, myeloperoxidase (MPO) in green, and nuclei (DAPI) in blue. Scale bar = 50 μm.

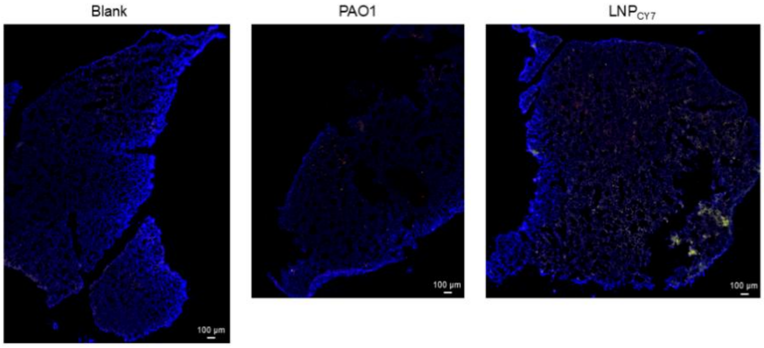


Supplementary Figure S9. Raw fluorescence images of the lungs from PAO-1 infected mice, acquired by panoramic scanning on the Mica Microhub (Leica Microsystems (Shanghai) Trading Co., Ltd.). Citrullinated histone H3 (Cit-H3) is shown in red, myeloperoxidase (MPO)

in green, and nuclei (DAPI) in blue. Scale bar = 100 μm.

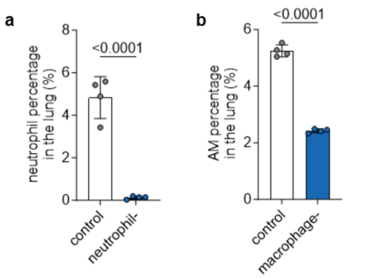


Supplementary Figure S10. Neutrophils or macrophage depletion confirmed by flow cytometry. a) Neutrophils were depleted by tail vein administration of 150 μg Ultra-LEAF™ anti-Ly6G antibody. Flow cytometric analysis confirmed efficient depletion of neutrophils in the lungs at 24 h post-treatment (n = 4). b) Alveolar macrophages (AMs) were depleted by intraperitoneal injection of 200 μL clodronate liposomes (5 mg/mL). Flow cytometric analysis confirmed efficient depletion in the lungs at 24 h post-treatment (n = 4). Data are presented as mean ± s.d. Statistical analyses were performed using unpaired t-test in Prism 10.

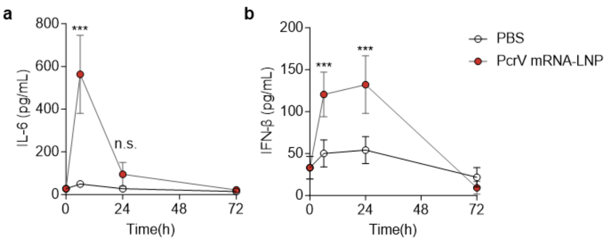


Supplementary Figure S11. IL-6 (a) and IFN-β (b) levels in BALF of vaccinated mice at indicated time points post-vaccination measured by ELISA (n=5). Data are presented as mean ± s.d. Statistical analyses were performed using two-way ANOVA in Prism 10. *** indicates $p < 0.001$; n.s. indicates no significant difference.

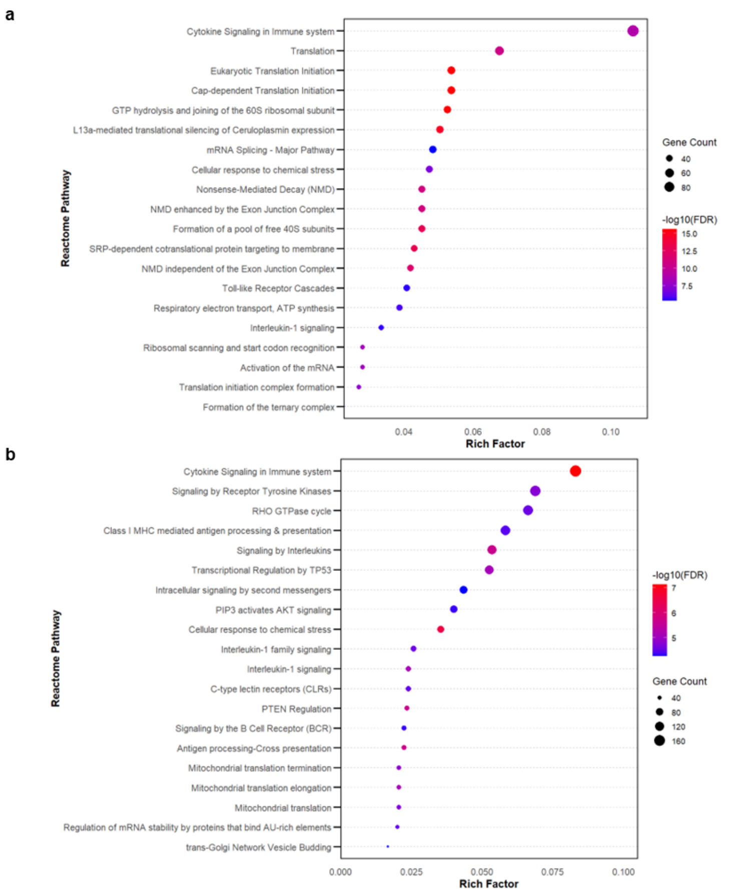


Supplementary Figure S12. Reactome analysis of neutrophils (a) and macrophages (b) 72 hours post pulmonary mRNA-LNP vaccination.

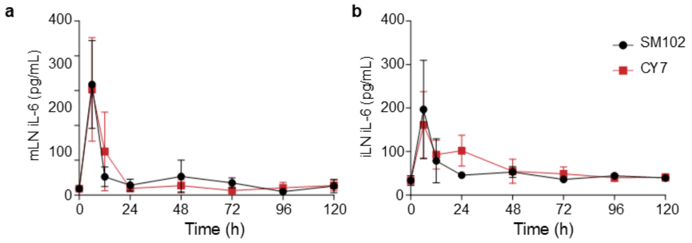


Supplementary Figure S13. IL-6 levels in lymph nodes following immunization with rPcrV-adjuvanted eLNPs. BALB/c mice received a single immunization with 2 µg recombinant PcrV adjuvanted with $eLNP_{SM102}$ or $eLNP_{CY7}$ via intramuscular or intratracheal administration. IL-6 levels were measured by ELISA in lysates of mediastinal (a) or iliac lymph nodes (b) at indicated time points post-immunization. Data are presented as mean ± s.d. (n = 3).

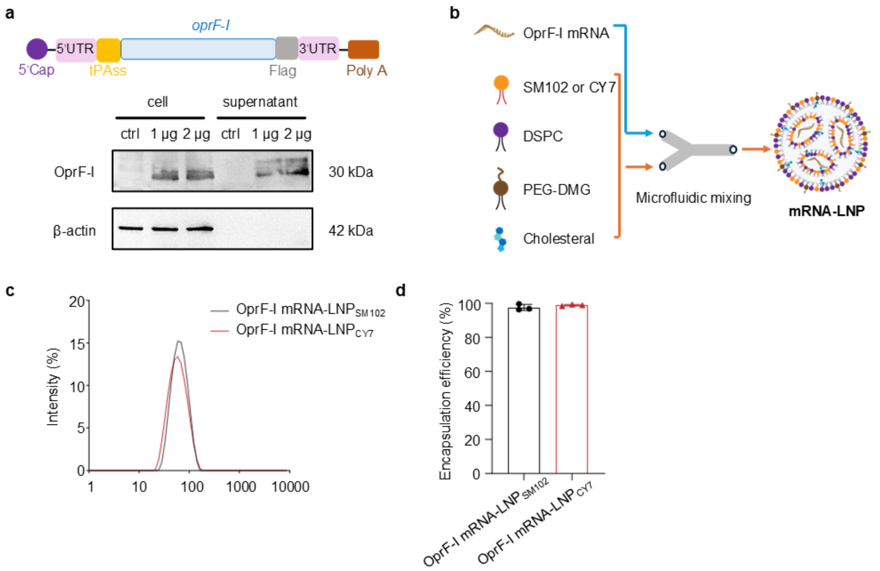


Supplementary Figure S14. OprF-I mRNA-LNP fabrication and characterization. a) Schematic representation of the OprF-I mRNA construct, consisting of a 5′cap, 5′untranslated region (5′UTR), signal peptide, coding sequence of OprF-I, 3′UTR, and a poly(A) tail. Western blot analysis of OprF-I protein expression and secretion 24 hours after transfection of HEK-293T cells with 1 μg or 2 μg OprF-I mRNA. The results indicate that the protein was secreted into the culture supernatant. b) Schematic representing the preparation of OprF-I mRNA-LNP. c) Particle size measured by dynamic light scattering (DLS). d) Encapsulation efficiency of OprF-I mRNA-LNPs determined by RiboGreen assay (n = 3).

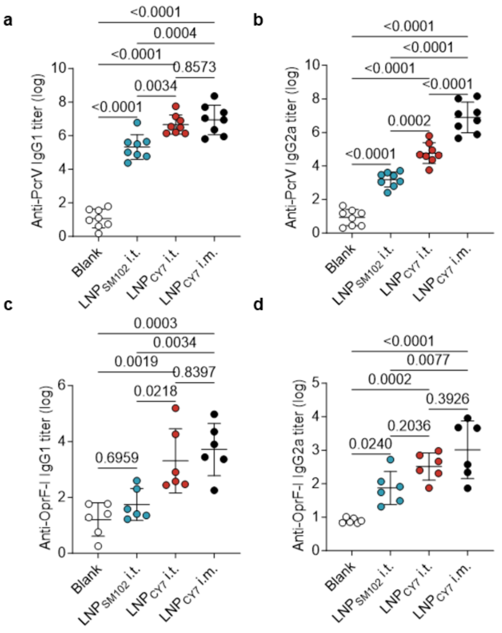


Supplementary Figure S15. Serum IgG1 and IgG2a titers against PcrV and OprF-I in immunized mice. Anti-PcrV IgG1 (a), Anti-PcrV IgG2a (b), anti-OprF-I IgG1 (c) and anti-

OprF-I IgG2a (d) antibody titers were measured in the sera of immunized mice on day 28 post-immunization (n = 6-8). Data are presented as mean ± s.d. Statistical analyses were performed using one-way ANOVA in Prism 10.

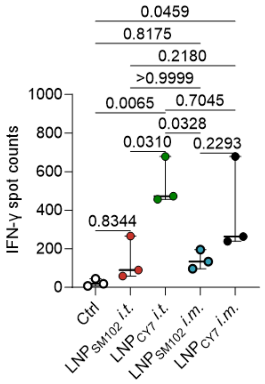


Supplementary Figure S16. Quantification of spot-forming units indicating IFN-γ-producing T cells in immunized mice. Spots were counted and analyzed using an automated ELISpot reader (n = 3). Data represent mean ± s.d. Statistical analyses were performed using one-way ANOVA in Prism 10.

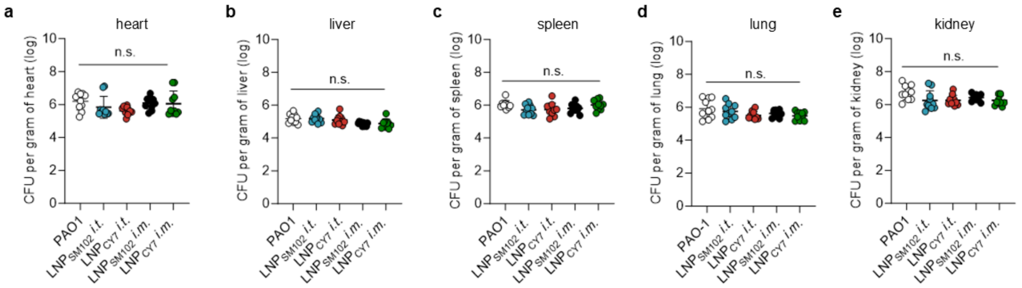


Supplementary Figure S17. Control efficacy study using Luc mRNA-LNPs formulated with either $LNP_{CY7}$ or $LNP_{SM102}$. BALB/c mice were immunized via i.t. or i.m. routes with 2 μg Luc mRNA-LNPs, followed by intravenous challenge with PAO1. Bacterial burdens in different organs were quantified 24 hours post-infection (n = 10 per group). No significant protection was observed for Luc mRNA-LNPs regardless of LNP type or administration route. Data are presented as mean ± s.d., and statistical significance was determined using one-way ANOVA with Tukey's multiple comparisons test.

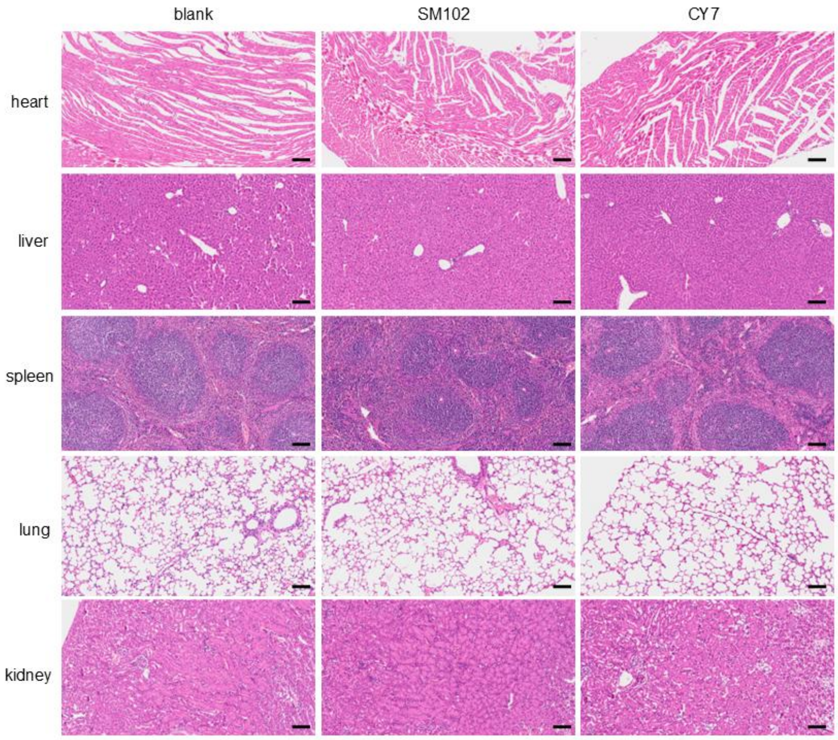


Supplementary Figure S18. Representative HE-stained sections of the heart, liver, spleen, lungs, and kidneys collected 24 h after intratracheal immunization. No observable pathological alterations were detected, indicating no evident organ toxicity. Scale bar, 100 μm.

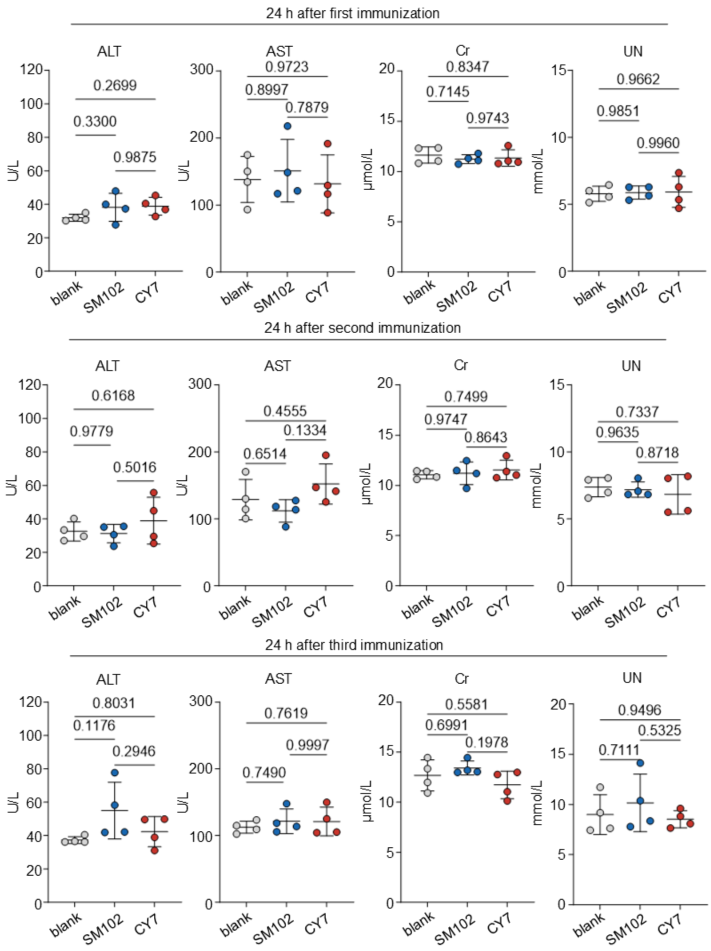


Supplementary Figure S19. Biochemical analyses of blood ALT, AST, Cr, and BUN levels at 24 h after each intratracheal administration showed that hepatic and renal functions remained comparable to those of the blank control group, indicating no evident systemic toxicity following pulmonary immunization. Data are presented as mean ± s.d (n=4). Statistical analyses were performed using one-way ANOVA in Prism 10.

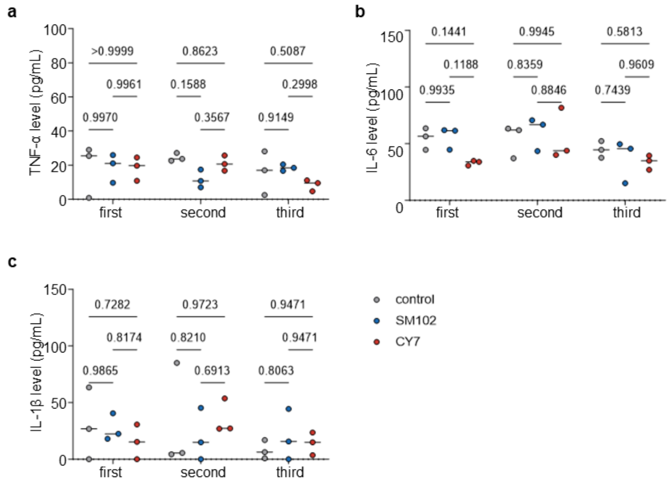


Supplementary Figure S20. Systemic cytokine levels following each vaccination. Serum concentrations of TNF-α (a), IL-6 (b), and IL-1β (c) measured 24 hours after intratracheal administration of mRNA-LNP$_{SM102}$ and mRNA-LNP$_{CY7}$. Data are presented as mean ± s.d (n=3). Statistical analyses were performed using two-way ANOVA in Prism 10.
